# Resource sharing by outer membrane vesicles from a citrus pathogen

**DOI:** 10.1101/2021.04.26.441564

**Authors:** Gabriel G. Araujo, Matheus M. Conforte, Aline D. da Purificação, Iris Todeschini, Edgar E. Llontop, Claudia B. Angeli, Alex Inague, Marcos Y. Yoshinaga, Robson F. de Souza, Rodrigo Papai, Maciel S. Luz, Sayuri Miyamoto, Giuseppe Palmisano, Chuck S. Farah, Cristiane R. Guzzo

## Abstract

The causative agent of citrus canker disease, *Xanthomonas citri* pv. *citri*, was found to produce copious amounts of outer membrane vesicles (OMVs), frequently forming long membranous tubes under different culture conditions. Lipidomic analysis revealed significant differences in lipid composition between purified vesicles in relation to whole cells. The results suggest an enrichment in saturated cardiolipins and a decrease in unsaturated lipids in the OMV samples, possibly granting them a more rigid structure while allowing their high degree of curvature caused by their small diameters. The vesicles’ proteome was found to be significantly enriched in TonB-dependent receptors related to the acquisition of different nutrients. These proteins are known to transport siderophores, which were evidenced to be present in purified *X. citri* OMVs, along with essential metals including iron, zinc, and manganese quantified by elemental analysis. The availability of vesicle-associated nutrients to be incorporated by cells was demonstrated by the use of OMVs as the sole carbon source for bacterial growth. At last, the vesicles also presented esterase and protease activities, which have been associated with virulence in phytopathogens. These evidences point that *X. citri* cells can use OMVs to share resources within microbial communities, which has potential implications for microbial interactions and plant colonization, affecting their survival and persistence on the host and in the environment.

**Importance:** The shedding of outer membrane vesicles appears to be universal in Gram-negative bacteria and effectively constitutes a unique secretion pathway for diverse molecules and proteins. To study their possible functions in the citrus pathogen *Xanthomonas citri*, purified vesicles from this bacterium were studied by omics and functional approaches. Nutrient transporters were found associated to these structures, which were evidenced to contain siderophores and essential metals. The availability of these nutrients to be incorporated by cells was then demonstrated by showing that purified vesicles can be used as sole carbon sources for microbial growth. Additionally, the samples also presented esterase and protease activities which can contribute to the release of substrates from plant host tissues. These observations help to establish the developing idea of vesicles as shared bacterial resources which can participate in shaping host-associated microbial communities in contrast to other interactions such as bacterial competition.

## Introduction

The production of outer membrane vesicles (OMVs) is known to be extremely common to Gram-negative bacteria and has been specially explored in pathogens due to their association to virulence factors (Schwechheimer and Kuehn, 2015; Toyofuku et al., 2019). Less commonly described structures are outer membrane tubes, also named tube-shaped membranous structures, nanotubes, nanowires and nanopods in different organisms. The tubes are considered to be a specialized form of OMVs, which assemble in the form of chains or completely fused to one another (Pirbadian et al., 2014; Remis et al., 2014; Pirbadian et al., 2015; Fischer et al., 2019; Toyofuku et al., 2019). These tubes seem to have the potential to bridge cell surfaces at long ranges, but their exact function, if at all dependent on their elongated shape, is still unclear on most cases and varies between different organisms.

*Myxococcus xanthus* outer membrane tubes are some of the most studied of these structures, forming a widespread network between the cells within biofilms that were proposed to promote coordination for these bacteria’s notorious social behaviors by serving as a transport medium for proteins and other molecules (Remis et al., 2014). Nevertheless, simply the presence of the tubes may not be sufficient for such activities since specific factors were found to be necessary to allow effective molecular exchanges through outer membrane connections. Namely, the proteins TraA and TraB were identified by genetic screening to be required for transferring outer membrane proteins by direct contacts between cells, while not affecting the production of tubes (Dey and Wall, 2014; Cao and Wall, 2019). In the zoonotic pathogen *Francisella novicida* which causes tularemia disease, virulence factors were detected in its OMVs and outer membrane tubes, which interestingly always appear to be of a continuous, non-segmented type. Interaction with host cells led to increased expression of the tubes, suggesting a role of these structures in the infection process (McCaig et al., 2013; Sampath et al., 2018). In *Vibrio vulnificus*, OMVs carry the virulence factor cytolysin–hemolysin VvhA (Kim et al., 2010), while its segmented tubes seem to exist only transiently as intermediates within the capsule of this opportunistic pathogen (Hampton et al., 2017). Somewhat in contrast to these examples, the outer membrane tubes of *Shewanella oneidensis* seem play a much clearer role in the biology of this organism. These membranous extensions form “nanowires” from which components of the electron transport chain of this metal-reducing bacterium can reach extracellular mineral electron acceptors (Pirbadian et al., 2014, 2015).

Studies with other environmental bacteria also revealed other possible implications of these structures on cell metabolism. In a marine *Flavobacterium* sp., OMV chains were proposed to serve as an extension of the cell surface for the degradation and incorporation of substrates (Fischer et al., 2019). OMVs of polycyclic aromatic hydrocarbon-degrading *Delftia* sp. Cs1-4 were found to be contained within tubular “nanopods” surrounded by a surface layer protein, NpdA, the production of which was stimulated by growth on phenanthrene. The presence of NpdA and the formation of an encasing structure for OMV tubes seem to be a characteristic distributed within the *Comamonadaceae* family (Shetty et al., 2011).

The examples presented above represent some of the exploration done on the relatively few, but nonetheless diverse bacteria identified that assemble extracellular tubular-shaped structures from their outer membrane. Nevertheless, OMVs are most commonly found not as chains but as free entities, which are produced by Gram-negative bacteria in different environments, such as biofilms, planktonic cultures, and within hosts (Hellman et al., 2000; Biller et al., 2014; Hickey et al., 2015). More generally speaking, extracellular membrane vesicles are also commonly produced by Gram-positive bacteria, archaea, and by eukaryotic cells (Schwechheimer and Kuehn, 2015).

Due to them being an effective way for microbial cells to release the most diverse compounds, OMV production can be used as a secretion mechanism and thus have been called the “type zero secretion system” (Schwechheimer and Kuehn, 2015; Guerrero-Mandujano et al., 2017; Toyofuku et al., 2019). Differently from other bacterial secretion systems, OMVs require a remodeling of the Gram-negative envelope to release vesicles made of outer membrane constituents with a periplasmic lumen. Therefore, different bacterial envelope crosslinks and non-covalent interactions between proteins located in the membrane that interact with the cell wall must be broken during the secretion of OMVs (Schwechheimer and Kuehn, 2015). Details of this process are still unclear, as well as if there is any generalized protein system actively involved in OMV biogenesis. Another process that is still not well understood is cargo selection, if proteins and chemical compounds can be directed into OMVs by the cell and secreted to the extracellular medium (Lappann et al., 2013; Elhenawy et al., 2014). In some bacteria, OMV synthesis can be triggered under stress conditions, such as antibiotic treatments that activate SOS response and under oxidative stress (McBroom and Kuehn, 2007; Maredia et al., 2012; Macdonald and Kuehn, 2013; Schwechheimer and Kuehn, 2013). Under these situations, OMVs may serve as a way to remove potentially harmful compounds, such as misfolded proteins.

OMVs may promote the acquisition of nutrients and essential ions such as iron and zinc in bacterial communities and during host colonization (Evans et al., 2012; Toledo et al., 2012; Biller et al., 2014; Schwechheimer and Kuehn, 2015). The role of OMVs in nutrition has been suggested for different bacteria, such as *M. xanthus*, the cyanobacterium *Prochlorococcus* sp., *Borrelia burgdorferi, Neisseria meningitidis, Porphyromonas gingivalis, Moraxella catarrhalis*, and for cytoplasmic membrane vesicles of *Mycobacterium tuberculosis* (Aebi et al., 1996; Evans et al., 2012; Toledo et al., 2012; Lappann et al., 2013; Biller et al., 2014). It is not clear if OMVs have a universal role for nutrient acquisition, but in some cases they have been suggested to act as public goods that benefit the producer cells as well as other bacteria from the community that can absorb them or use the products released by the action of enzymes located in the OMVs (Evans et al., 2012; Elhenawy et al., 2014; Schwechheimer and Kuehn, 2015). An example is the relationship between bacteria found in the gut microbiota. OMVs produced by *Bacteroides* species carry hydrolases and polysaccharide lyases which can be used by bacteria that do not produce these enzymes to metabolize polysaccharides as nutrient sources in a mutualistic interaction (Rakoff-Nahoum et al., 2014).

Few studies have focused in the OMVs of phytopathogens. Still, research on this topic has revealed that, similarly to their animal-colonizing counterparts, bacteria that inflict diseases on plants were found to produce vesicles loaded with virulence-associated proteins and are capable of inducing immune responses on their hosts (Sidhu et al., 2008; Solé et al., 2015; Bahar et al., 2016; Nascimento et al., 2016; Katsir and Bahar, 2017; Feitosa-Junior et al., 2019). These observations include *Xanthomonas* species and the closely related plant pathogen *Xylella fastidiosa*.

Strains from the genus *Xanthomonas*, known to cause diseases in a number of plant hosts, frequently contain most of the traditional bacterial macromolecular secretion systems named type I to VI (Büttner and Bonas, 2010; Alvarez-Martinez et al., 2021). OMVs, however, are comparatively much less studied than these other systems in these bacteria. Thus, this work focuses on unveiling the composition and possible roles of vesicles from one such phytopathogen, the causative agent of citrus canker disease, *Xanthomonas citri* pv. *citri* strain 306 (*X. citri*). Long extracellular appendages composed of OMVs were identified under different culture conditions, and the purified OMVs were investigated by elemental analysis, proteomic and lipidomic techniques, as well as by functional approaches. The vesicles were found to be potential vehicles of nutrients and essential ions available for incorporation by bacterial cells. This function, in association with the esterase and protease activities observed in the purified *X. citri* OMVs, may possibly aid in the microbial colonization of the plant host and contribute to disease establishment.

## Results and Discussion

### *Visualization of* X. citri *outer membrane vesicles and tubes by negative stain TEM*

Negative stain transmission electron microscopy (TEM) revealed the presence of tubular extensions from *X. citri* cells grown in plates of different culture media, identified as outer membrane tubes (**Fig. 1**). Upon closer inspection, the tubes were found to be formed from vesicle chains, occasionally with a well-defined segmentation but frequently presenting nearly indistinguishable boundaries between links, seeming almost continuous. The size of the tubes ranged from short segments up to a few micrometers in length. Surrounding the cells in all conditions tested, a multitude of outer membrane vesicles (OMVs) was also present (**Fig. 1**). For most of their extension, the tubes appear to be composed of vesicles with a more homogeneous diameter (58-74 nm) than the isolated OMVs. Each tube seem to possess larger vesicles at its tips (88-103 nm), and some longer tubes appear to be formed by segments connected by these larger subunits.

**Fig. 1.**
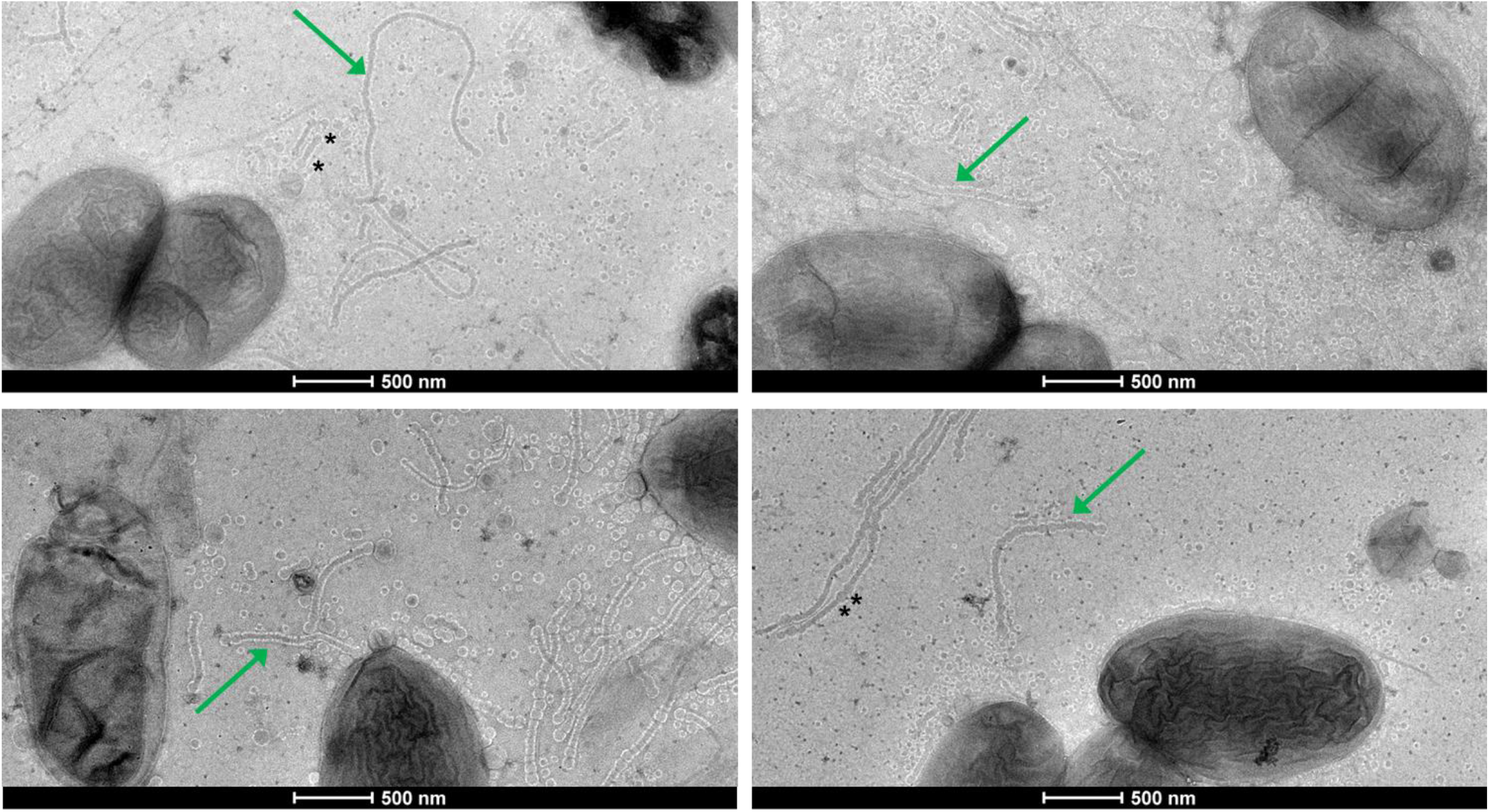
Outer membrane tubes and vesicles from *X. citri*. Cells were grown in SB with agar 0.6% and imaged by negative stain TEM. The green arrows point to examples of the outer membrane tubes that can be seen in the images. Asterisks (*) indicate some occurrences of larger vesicles at the tips or within tubes.

Amongst the different culture media tested, Silva–Buddenhagen (SB) plates (Ou, 1985) seemingly produced the largest amount of tubes and vesicles, and thus were used for further experiments. The agar percentage (0.6 – 1.5%) seemed to not significantly affect the production of tubes, but these structures seemed to be rarer when the cells were grown in liquid medium (**Fig. S1**).

### Purification of outer membrane vesicles combined with lipidomics and proteomics analyses

Pure, cell-free, OMVs could be purified from cultures grown in SB plates by filtration and density gradient centrifugation generating a clear yellow suspension (**Fig. 2A**). The tubes did not appear in the final preparations, being either lost during the process or disassembling from the manipulation (**Fig. 2B**). The purity of the OMV preparations was confirmed by negative stain TEM and absence of growth from contaminating cells. In addition to that, dynamic light scattering (DLS) was employed to measure their diameter distribution. The vesicles were determined to be monodispersed, with sizes ranging from about 40 to 150 nm, with a peak at around 75 nm (**Fig. 2C**), well within previous descriptions for OMVs (Schwechheimer and Kuehn, 2015). The purified samples were then subjected to different analytical procedures to reveal their molecular composition.

**Fig. 2.**
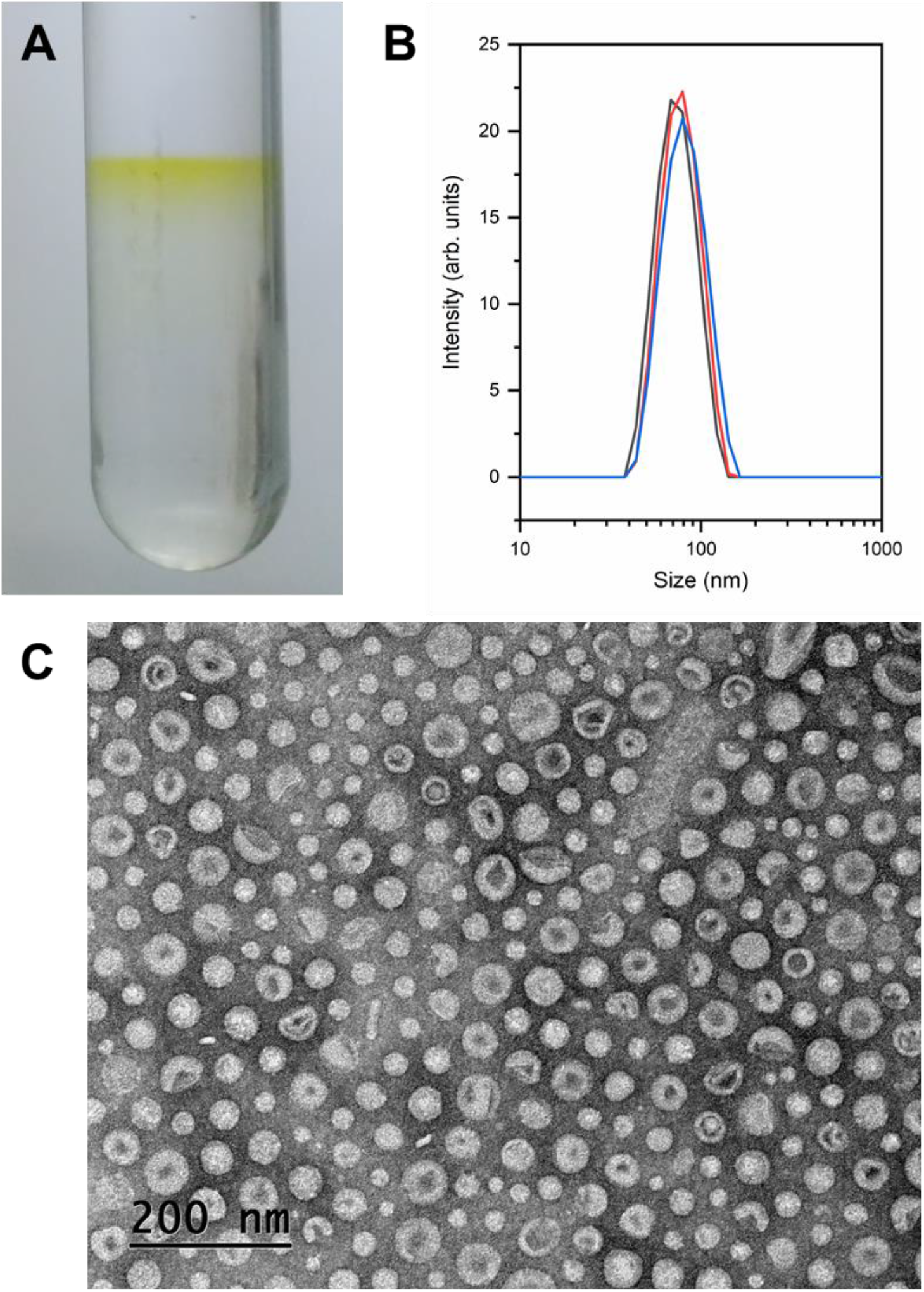
Purification and characterization of the *X. citri* outer membrane vesicles (OMVs). Vesicles were retrieved as a yellow band from the density gradient centrifugation tubes (A). The vesicle size distribution was determined by DLS and observed to range from about 40 to 150 nm in diameter, with a peak near 75 nm (B). Observation by negative stain TEM confirmed the purification of the OMVs, allowing the evaluation of their size, morphology, and lack of contaminating cells (C).

Lipidomic analysis by liquid chromatography-tandem mass spectrometry (LC-MS/MS) of pure OMVs, partially purified OMV preparations (“OMV-enriched” samples, in which the cells were removed by filtration but not submitted to the density gradient centrifugation step), and whole *X. citri* cells revealed substantial differences between the samples. Sixty-six different lipids were identified, divided into 6 subclasses: cardiolipins (CL), free fatty acids (FFA), phosphatidylcholine (PC), phosphatidylethanolamine (PE), phosphatidylglycerol (PG), and methylated-phosphatidylserine (PS-Me) (**Fig. 3A**). CL, a type of diphosphatidylglycerol lipid, was the most diverse and abundant lipid subclass in all samples (**Fig. 3B**). The main difference observed was that, in relation to whole cells, pure OMVs appeared to be enriched in CL and relatively impoverished in PG (the biosynthetic precursor of CL). Free fatty acids were highly prevalent, likely reflecting their important role as common metabolic intermediates. It is important to note that the main components of the outer leaflet of bacterial outer membranes, lipopolysaccharides (LPS), were not evaluated in this analysis due to their relatively hydrophilic nature, making them too polar to be extracted along the other lipids.

**Fig. 3.**
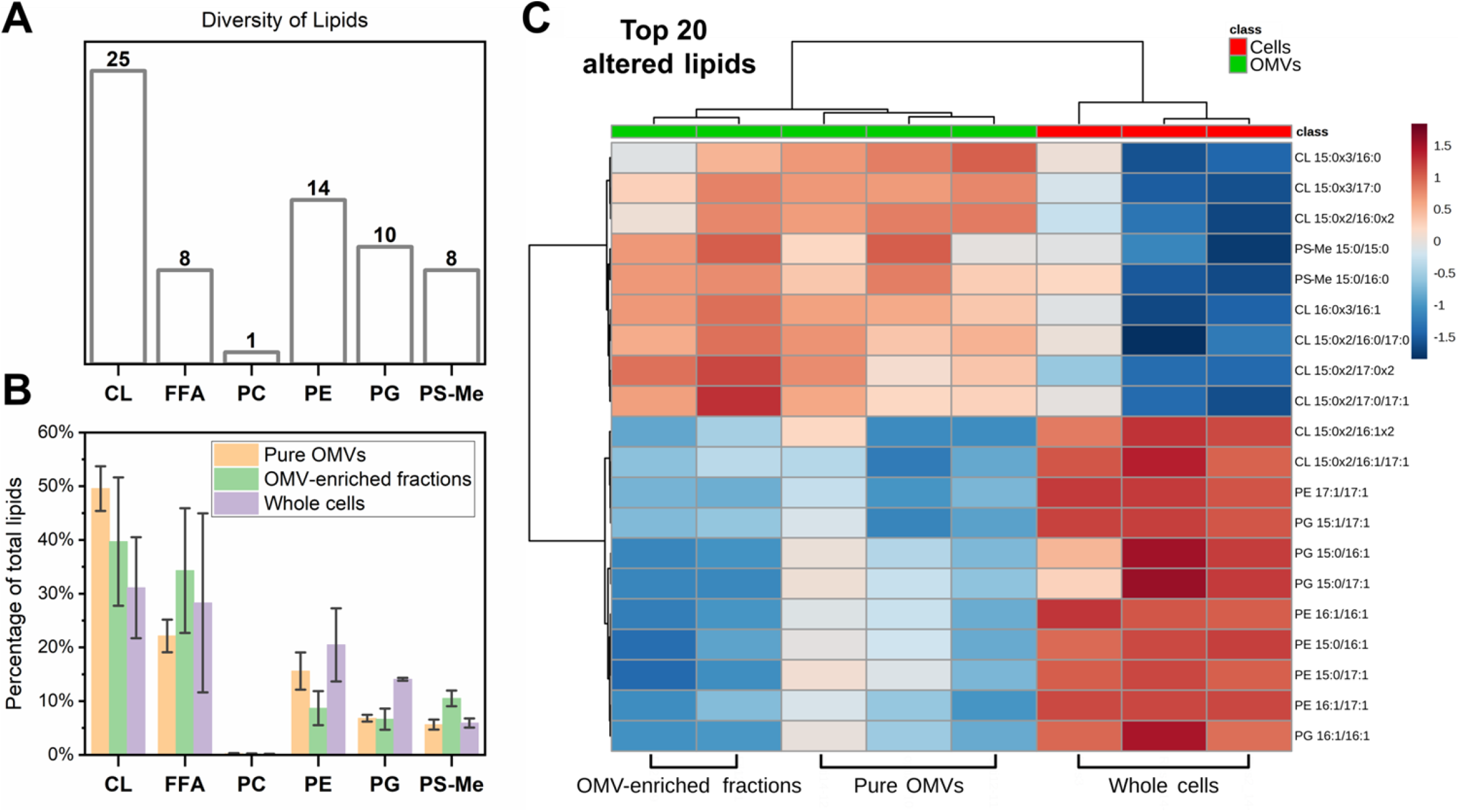
Lipidomic analysis of *X. citri* whole cells and OMVs. A total of 66 different lipids were identified in the samples, divided into 6 subclasses: cardiolipin (CL), free fatty acids (FFA), phosphatidylcholine (PC), phosphatidylethanolamine (PE), phosphatidylglycerol (PG), methylated-phosphatidylserine (PS-Me), shown in panel A. The proportion of each lipid subclass varied between the different samples: Pure OMVs, OMV-enriched fractions (partially purified), and whole cells (B). The 20 most altered lipids between the different samples (identified from a volcano plot analysis, fold-change > 1.5, p < 0.05 evaluated by FDR-adjusted t-test, **Fig. S2**) were clustered in a heatmap (according to one-way ANOVA), revealing the vesicles are enriched in saturated cardiolipins in comparison to the cells, while being relatively impoverished in a number of different unsaturated lipid species (C). The notation used to represent the lipids from the different subclasses gives the number of carbon atoms and of double bonds separated by a colon for each acyl chain, which in turn are separated by a slash.

A volcano plot analysis revealed 20 altered lipids between OMV-containing and whole cell samples, all presenting significant (p<0.05 ; FDR-adjusted t-test) fold changes values above 1.5 (**Fig. S2**). In the heatmap distribution for these altered lipids, according to one-way ANOVA, each sample type clustered with its replicates, with the OMVs (partially or completely purified) grouping separately from whole cells (**Fig. 3C**). Interestingly, it could be observed that the OMVs had relatively increased amounts of several CL species linked to saturated fatty acids and decreased quantities of phospholipids (including CL) linked to unsaturated fatty acids when compared to the whole cells.

The cone-shaped lipid CL is known to localize to negative curvature regions on membranes (Renner and Weibel, 2011; Beltran-Heredia et al., 2019), such as in the inner leaflet of *X. citri* OMVs, which present small diameters and are thus highly curved structures. Additionally, the relatively higher saturation of the CL-linked chains in the vesicles may grant the OMVs with more membrane rigidity (Tashiro et al., 2011). CL has been described as organizing into microdomains where CL-interacting proteins localize (Sorice et al., 2009; Planas-Iglesias et al., 2015; Lin and Weibel, 2016). In this manner, protein affinity for these lipids could contribute cargo sorting into *X. citri* OMVs.

Nanoflow liquid chromatography-tandem mass spectrometry (nLC-MS/MS) was used for the proteomic analysis of two replicates of purified OMV suspensions using in-solution digestion. Parallel to that, four bands of OMV proteins separated in a SDS-PAGE gel were used for a gel electrophoresis liquid chromatography (GeLC) approach using in-gel digestion (**Fig. 4A**). The data from the gel band samples were pooled and quantitatively compared to the two replicates of the in-solution digestion. A total of 698 proteins were identified with at least one peptide, with 561 proteins presenting two or more peptides (**Data Set S1** and **Data Set S2**). Using their iBAQ (intensity based absolute quantification) values, the top 100 most abundant proteins from each sample were selected and compared (**Fig. 4B**). While the in-solution duplicates presented a large overlap, sharing 86 of their top 100 proteins, the GeLC approach (gel bands samples) revealed the most distinct profile, with 49 of their most abundant proteins being unique to its set.

**Fig. 4.**
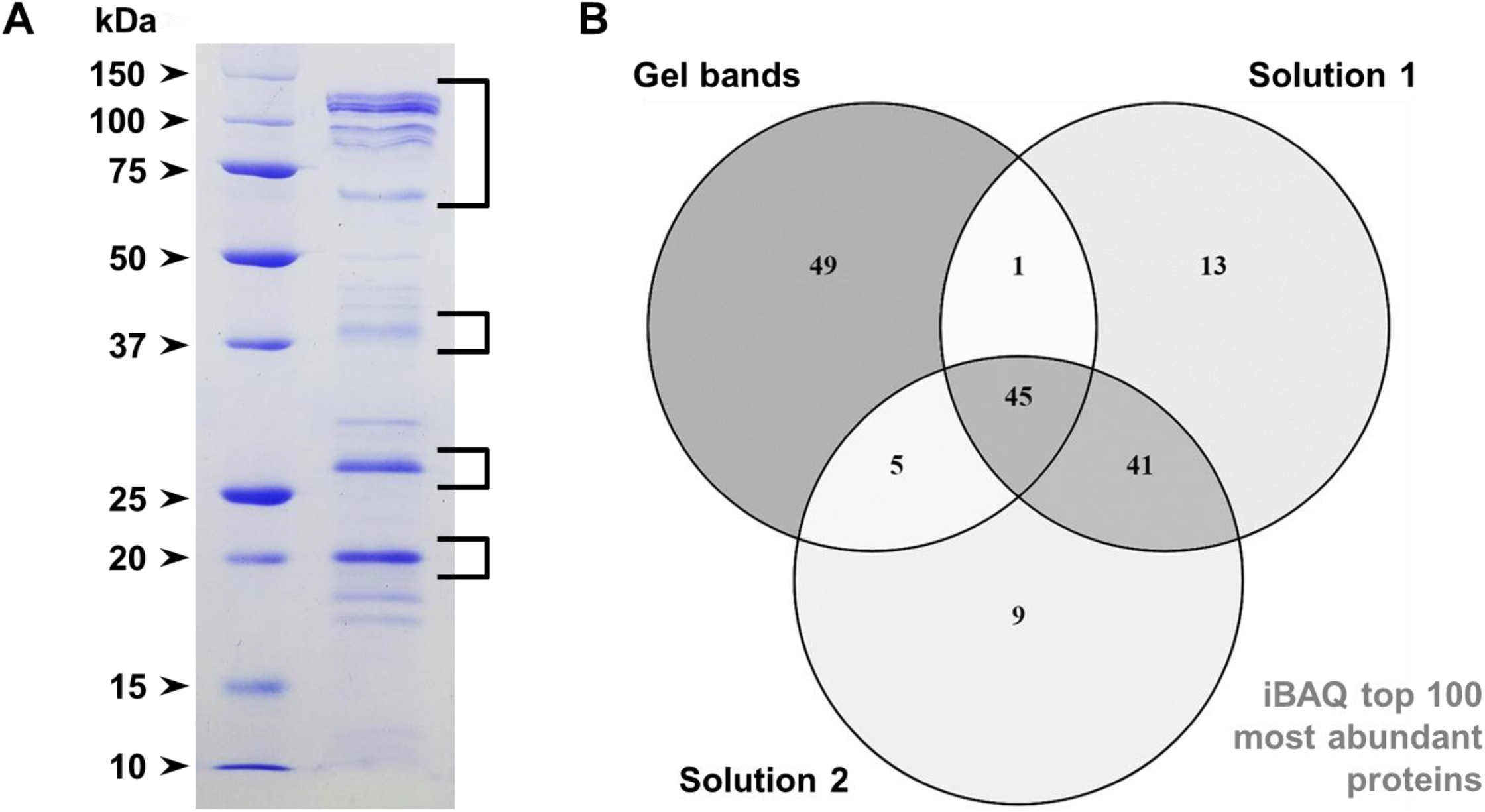
Proteomic analysis of *X. citri* OMV samples. Panel A shows characteristic protein bands associated with the purified OMVs that could be observed in 15% Tris-Glycine SDS-PAGE gels. Four regions containing the main bands (square brackets) were processed by in-gel digestion for proteomic analyses; their data were combined (“gel bands”) and compared to two samples of pure OMV suspensions processed by in-solution digestion (“solution 1” and “solution 2”). Panel B presents a Venn diagram displaying the intersection of the top 100 most abundant proteins for each sample determined by their iBAQ values.

The grouping of the top 100 non-redundant proteins with the highest iBAQ values for each sample yielded a list of 163 different proteins (**Table S1**). Subcellular localization prediction with PSORTb, manually curated based on sequence annotations, pointed out that 42.3% of these sequences are expected to be outer membrane proteins and 12.3% are likely periplasmic (**Fig. 5A**). The presence of inner membrane and cytoplasmic components observed in the proteome of *X. citri* OMVs, including ribosomal proteins, is commonly reported in the literature but remains unexplained as to how these proteins might associate to OMVs (Schwechheimer and Kuehn, 2015; Sjöström et al., 2015; Toyofuku et al., 2019; Zwarycz et al., 2020). Additionally, a cellular location could not be predicted for 21.5% of the identified proteins. For a different view on protein localization, SignalP was used to predict the secretion mechanisms of the OMV proteins (**Fig. 5B**). Nearly half of them (49.7%) contained signal peptides and almost one-fifth (19%) were predicted lipoproteins. A large “other” category (29.4%) includes cytoplasmic components and other proteins with non-classical or unknown secretion mechanisms.

**Fig. 5.**
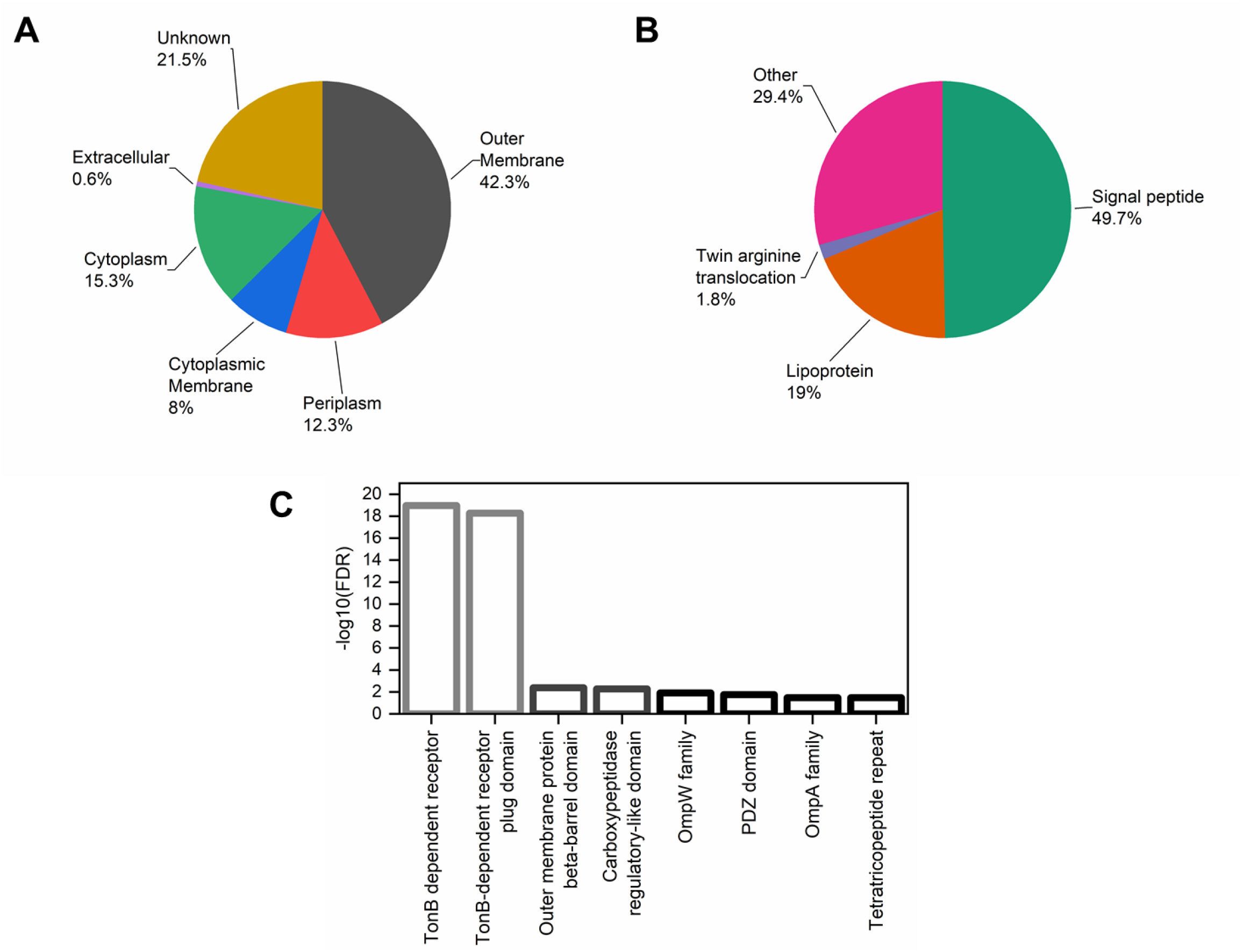
Subcellular localization and domain enrichment of the most abundant proteins identified in the purified *X. citri* OMV samples. Panel A presents the subcellular protein localization predicted by PSORTb, manually curated based on sequence annotations, while panel B shows their secretion mechanisms predicted by SignalP. Panel C displays the most significantly enriched Pfam domains found in the OMVs compared to the *X. citri* pv. *citri* 306 genome. The lowest false discovery rate (FDR), thus the highest -log10(FDR), was observed for TonB-dependent receptor domains (Pfam family PF00593). These analyses were performed with the combination of the top 100 proteins with the highest iBAQ values from the different samples analyzed by proteomics (gel bands, solution 1, solution 2), resulting in a list of 163 non-redundant proteins (**Table S1**).

From a functional perspective, proteins containing a TonB-dependent receptor domain were the most significantly enriched in the vesicles in comparison to the *X. citri* pv. *citri* 306 genome (**Fig. 5C**), as determined for Pfam annotations by the statistical enrichment analysis function of the STRING database (Franceschini et al., 2013). In accordance with that, the STRING analysis also identified a number of InterPro domains related to TonB-dependent receptors as the most significantly enriched in the samples (**Fig. S3**). In total, of the 163 most abundant proteins (**Table S1**), 31 were found to contain a “TonB-dependent receptor” Pfam domain (PF00593), the same set which contained a “TonB-dependent receptor-like, beta-barrel” InterPro domain (IPR000531). In a previous report, TonB-dependent receptors were found to compose the majority of the identified outer membrane proteins in OMVs from *Xanthomonas campestris* pv. *campestris* (Sidhu et al., 2008). These outer membrane receptors are known to transport a range of nutrients, including metal-binding compounds (particularly siderophores), nickel complexes, vitamin B_12_, and carbohydrates (Blanvillain et al., 2007; Krewulak and Vogel, 2011). Based on sequence annotations, the OMV proteome presents different types of TonB-dependent receptors which may bind diverse substrates (**Table S1**). These proteins are expected to remain able to bind to their specific ligands in the surface of the OMVs, though their internalization should not occur under these conditions since inner membrane components of this transport system are necessary to power substrate translocation (Krewulak and Vogel, 2011).

### OMVs as sources of nutrients and essential metals

Based on similar observations in relation to ion transporters in their proteomes, OMVs from different bacterial species have been suggested to be involved in metal acquisition (Schwechheimer and Kuehn, 2015). Given this abundance of TonB-dependent receptors in the *X. citri* OMVs, mainly associated with siderophore transport, chrome azurol S (CAS) agar plates (Schwyn and Neilands, 1987) were used as a qualitative assay to evidence the presence of this type of molecule associated with the purified vesicles. OMV suspensions added to the medium caused its discoloration, indicating the displacement of the iron in the blue-colored CAS complex by the putative high affinity siderophores present in the samples (**Fig. 6A**). It is interesting to note that the iron-scavenging role of siderophores for microbial growth can also be important in phytopathogens for interactions with the host plant, promoting virulence and potentially triggering immune responses (Aznar and Dellagi, 2015).

**Fig. 6.**
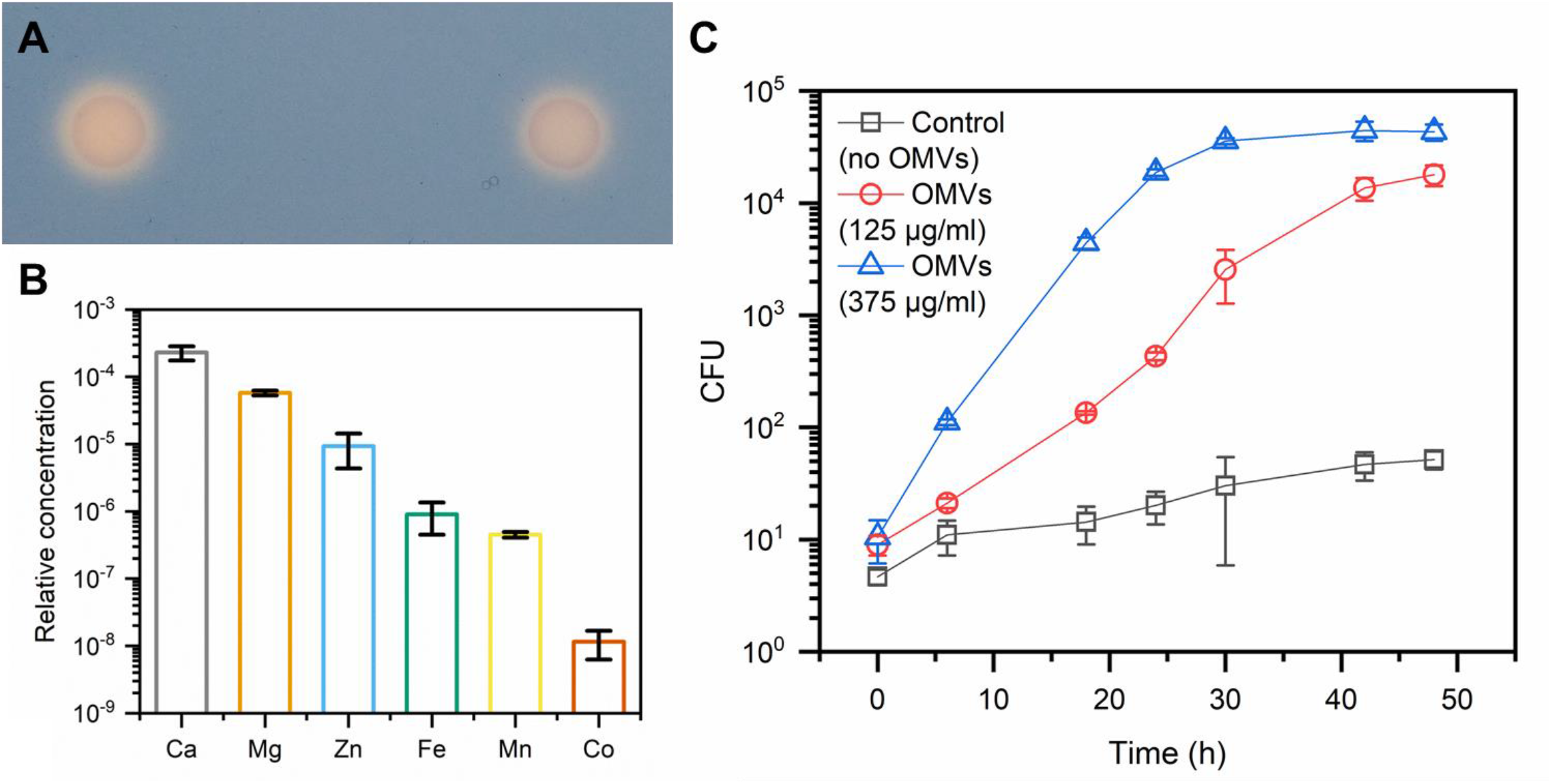
*X. citri* OMVs carry essential metals and can be incorporated by cells. Siderophores were potentially detected in the OMVs by discoloration of the medium in CAS plates where vesicles were applied (A). Elemental analysis of the OMVs revealed the presence of biologically important metals in the samples, including iron and zinc. The relative concentration (y-axis) was calculated by the ratio between the mass fraction values for each element and the carbon content. The oxidation state of each element was not determined. (B). *X. citri* can use OMVs as the sole carbon source for growth, indicating that the content of the vesicles is available for incorporation by cells (C). Different OMV concentrations, measured by their protein content, were added to tubes with M9 medium without other carbon source and a substantial increase in CFU was observed after incubation.

To further investigate the association of the OMVs with essential metals, their elemental composition was determined by triple quadrupole inductively coupled plasma-mass spectrometry (TQ ICP-MS) (**Fig. 6B**). **Table S2** presents the full results for all tested elements (C, Mg, S, Ca, Mn, Fe, Co, Ni, Cu, Zn, Br, Se, I, and Ba). The relative concentration of the elements in relation to carbon, reported as element-to-carbon ratios, was used as a comparative abundance value of the chemical elements in OMVs. The analysis confirmed the presence iron in the OMVs (899±450 ppb in relation to carbon), concurrent with the observed occurrence of receptors for iron-binding molecules in the vesicles and evidence for the presence of siderophores in the samples. Yet, iron was found at a smaller concentration than calcium (229±54 ppm) and magnesium (58±5 ppm), which are probably mostly bound to the LPS layer on the vesicles’ surface (Coughlin et al., 1983), thus explaining their relative abundance. Zinc (9±5 ppm) was another biologically important metal ion determined at substantial levels in the OMVs. It can be used as a cofactor for different enzymes, including for metallopeptidases known to contribute to the pathogenicity of some organisms (Hase and Finkelstein, 1993). In fact, a few such zinc-dependent metallopeptidases were identified in the OMVs (**Data Set S1** and **Data Set S2**), though their specific biological roles have not yet been defined. In addition to that, manganese (450±45 ppb) can also act as a cofactor in a number of different enzymes and was also quantified in the samples. At last, cobalt (12±5 ppb) was detected in the vesicles. This is interesting given that among the TonB-dependent receptors enriched in the OMVs (**Table S1**), at least one is annotated as specific for vitamin B_12_, a molecule which contains a coordinated cobalt ion. This protein, XAC3194, specifically contains a “TonB-dependent vitamin B_12_ transporter BtuB” InterPro Domain (IPR010101).

To test if the vesicles and the material associated to them are accessible to *X. citri* cells and can be utilized by them as nutrient sources, purified OMVs were tested as the sole carbon source for microbial growth. Substantial growth was observed for the samples where OMVs were added, with the highest vesicle protein concentration tested leading to a multiplication of about 1000-fold in colony-forming units (**Fig. 6C**), indicating that the macromolecules associated with the vesicles were being consumed by the bacteria. This confirms the ability of these structures and the material they carry to be incorporated and used by cells, strengthening the hypothesis that they can act as nutrient vehicles such as has been proposed for other bacteria (Aebi et al., 1996; Evans et al., 2012; Toledo et al., 2012; Lappann et al., 2013; Biller et al., 2014; Schwechheimer and Kuehn, 2015). The mechanism for this incorporation, however, remains unclear. It could be mediated by the degradation of the vesicles for the release of their contents in some manner, but fusion of the OMVs to the cells’ surfaces can also possibly be considered (Evans et al., 2012).

### Esterase and protease activity of OMVs

Additional functional assays with the purified *X. citri* OMVs revealed they present esterase activity. Qualitative assays on agar plates evidenced their capacity to cause the hydrolysis of the triglyceride tributyrin emulsified in the medium, generating a clear halo (**Fig. 7A**), as well as to release the fatty acids from molecules of Tween 20, leading to their precipitation with the calcium added to the plates (**Fig. 7B**). Further assays were performed in suspension with *p*-nitrophenyl butyrate (*p*NP-C4) and *p*-nitrophenyl octanoate (*p*NP-C8) as chromogenic substrates, adding controlled amounts of vesicles quantified by their protein content. Using *p*NP-C4, a clear trend could be observed of increasing OMV protein concentration leading to faster product release (**Fig. 7C**). The longer chain substrate *p*NP-C8 was also hydrolyzed, but there were no clear differences between the different quantities of added vesicles (**Fig. 7D**). This is probably due to the low solubility of *p*NP-C8 in the medium, thus becoming the limiting factor for the reaction. Nonetheless, with both *p*-nitrophenyl esters, a plateau seems to have been reached during the incubation with the OMVs, suggesting all the available substrate was consumed.

**Fig. 7.**
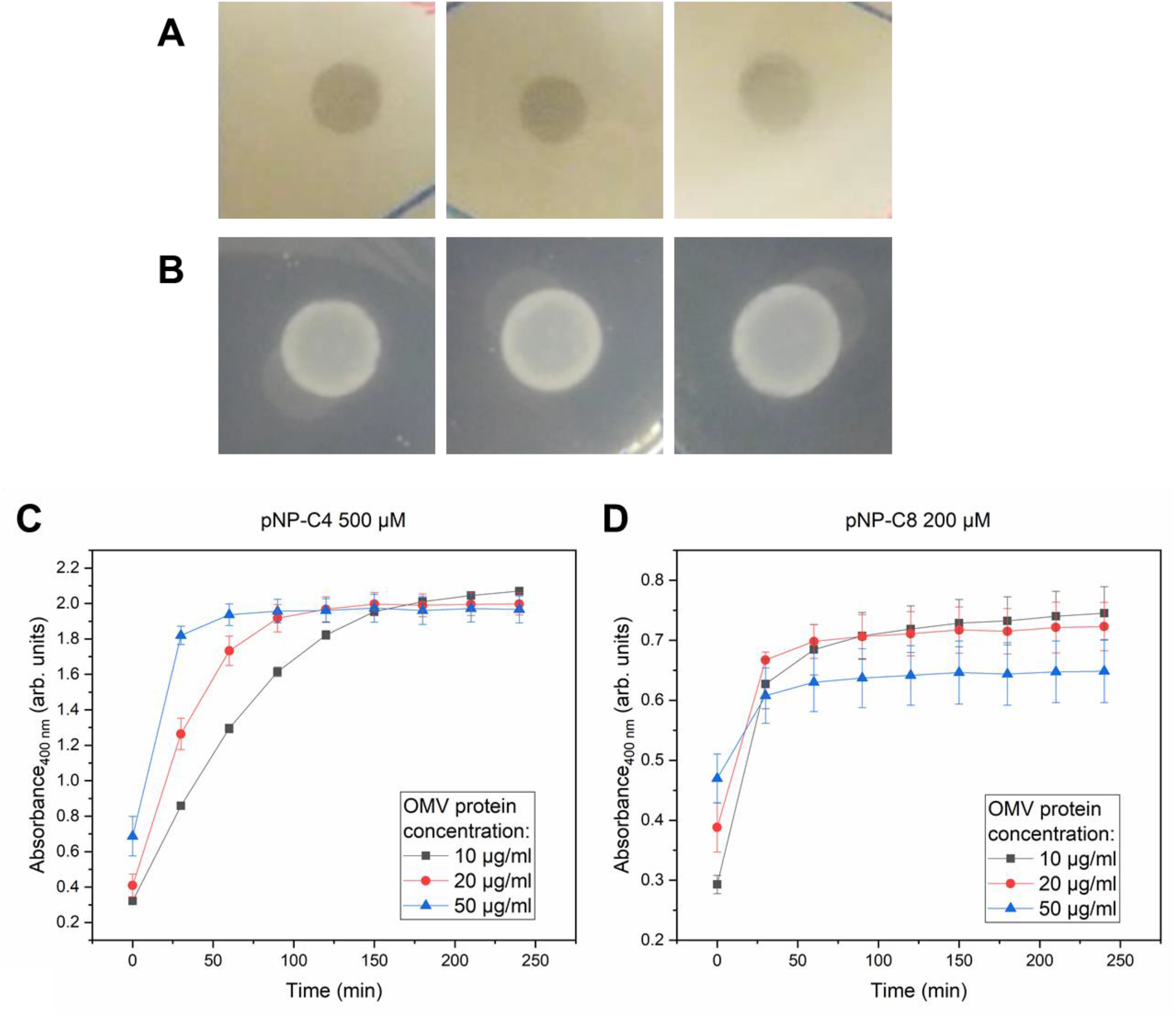
*X. citri* OMVs present esterase activity against a broad range of substrates. In qualitative esterase activity assays on agar plates, different purified OMV samples were able to create a clear halo in plates emulsified with the triglyceride tributirin (A) and to generate opaque white precipitates in plates containing Tween 20 and CaCl_2_ (B). These results indicate the hydrolysis of the respective substrates in the plates. Different OMV concentrations, measured by their total protein content, were able to hydrolyze *p*NP-C4 (panel C) and *p*NP-C8 (panel D) in colorimetric assays, indicated by the increase in absorbance at 400 nm during incubation.

The esterase activity associated with the OMVs measured for a broad range of substrates can possibly be attributed to the outer membrane esterase with an autotransporter domain XAC3300 (gene name *estA*) identified among the most abundant proteins in the proteome, though other undetected enzymes may be present. Esterases have been reported to contribute to the virulence of phytopathogens, playing roles such as aiding in the degradation of cutin, pectin, or xylan in plant host tissues (Fett et al., 1992; Aparna et al., 2009; Tamir-Ariel et al., 2012; Dejean et al., 2013; Nascimento et al., 2016; Tayi et al., 2016; Ueda et al., 2018), depending on their substrate preference. In *Xanthomonas oryzae* pv. *oryzae*, loss of function of the LipA esterase lead to loss of virulence on rice and to the inability to induce host defense responses (Aparna et al., 2009), while a LipA mutant of *Xanthomonas campestris* pv. *vesicatoria* induced less severe symptoms on tomato than the wild type (Tamir-Ariel et al., 2012). The LipA ortholog of the related plant pathogen *Xylella fastidiosa*, LesA, was found to be present in OMVs. This esterase was able to induce hypersensitive response-like symptoms in grapevine leafs, while a LesA mutant showed decreased virulence (Nascimento et al., 2016). At last, a LipA mutant of *X. citri* presented reduced symptoms when inoculated into citrus leaves (Assis et al., 2017). This particular protein (XAC0501), however, could not be identified in the *X. citri* OMV proteome (**Data Set S1** and **Data Set S2**) but other esterases could perform similar functions in the plant host.

Proteases are another class of hydrolases that have been associated to pathogenesis in plant-infecting microorganisms (Hou et al., 2018; Figaj et al., 2019). In the *X. citri* OMV samples, this enzymatic activity was identified utilizing a fluorescent casein substrate (**Fig. 8**), revealing yet another function connected to these structures. More substrate degradation was observed with the addition of increasing amounts of OMVs to the reactions, while a commercial EDTA-free protease inhibitor mix was able to substantially reduce activity (**Fig. 8**). In *X. campestris* pv. *campestris*, a protease-deficient mutant presented a substantial loss of pathogenicity in turnip leaves (Dow et al., 1990), whereas the XCV3671 protease of *X. campestris* pv. *vesicatoria* was determined to contribute to virulence in pepper plants and evidenced to be secreted in association with OMVs from this strain (Solé et al., 2015). Further research could show if similar enzymes, both proteases and other esterases, are important for *X. citri* infection and citrus canker development.

**Fig. 8.**
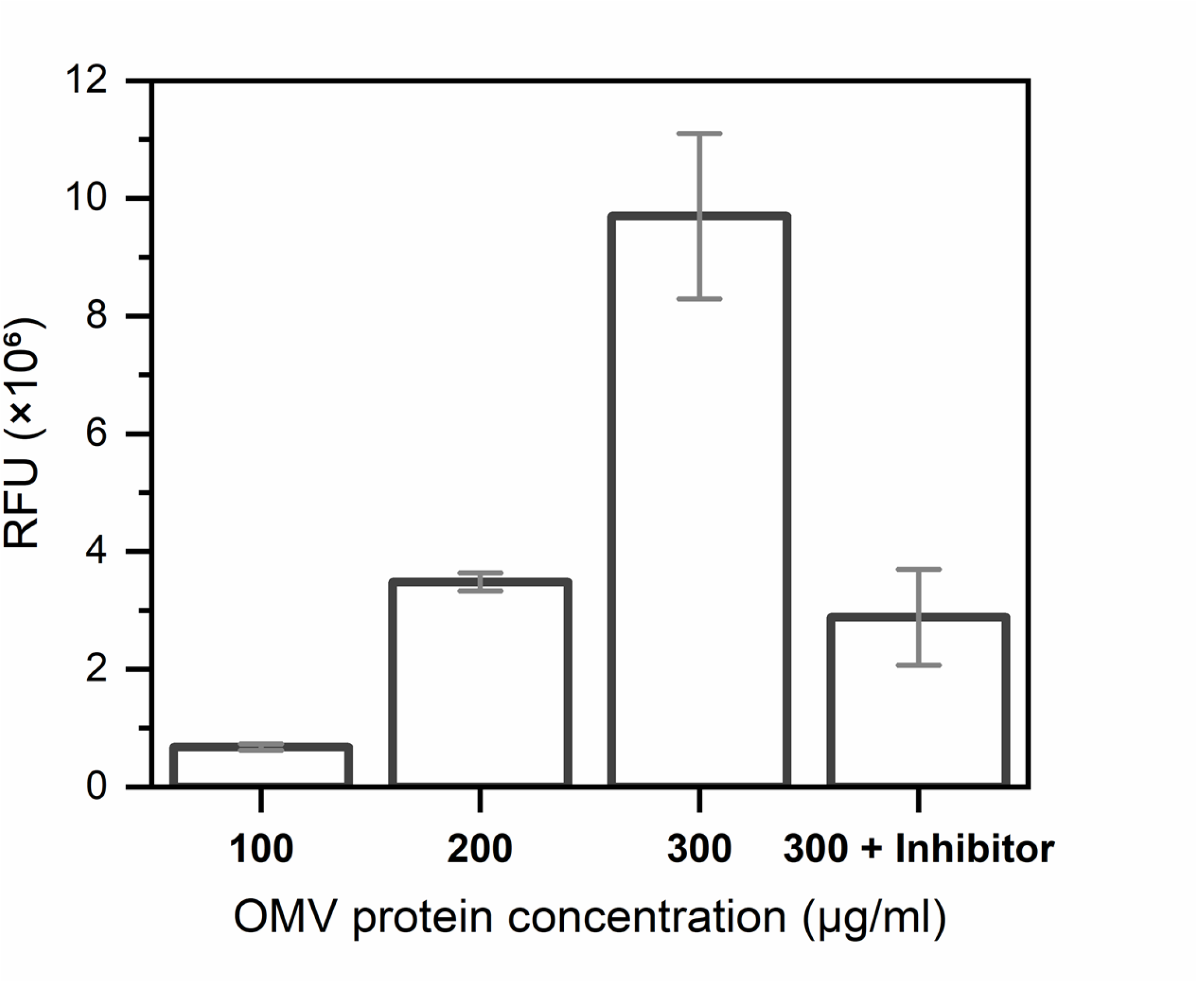
*X. citri* OMVs present protease activity. A protease fluorescent detection kit was used to detect the activity of purified OMVs at different concentrations, measured as relative fluorescence units (RFU). Addition of an EDTA-free protease inhibitor to samples with the highest OMV concentration tested lead to substantial decrease in the observed enzymatic activity. The fluorescence from blank (phosphate buffered saline) was subtracted from all samples.

Considering the identified enzymatic activities associated with the *X. citri* OMVs, as well as the presence of other hydrolases detected by proteomic analyses (**Table S1**), the vesicles may be an important resource in plant colonization and pathogenesis. The release of products from the degradation of macromolecules can be another manner in which the vesicles would be related to nutrient acquisition, acting as public goods for other *X. citri* cells and possibly the microbial community in general. These processes can facilitate the bacterial colonization of plant tissues and thus participate in disease development.

## Conclusion

*X. citri* cells express outer membrane tubes and vesicles carrying proteins, molecules, and ions that may benefit bacterial cells. The OMV lipid profile revealed their higher content of saturated cardiolipins with a relative impoverishment in unsaturated lipids. This might grant them more rigidity while maintaining the small diameter of the vesicles. The proteome of the vesicles revealed an abundance of transporters related to the uptake of nutrient molecules from the medium. This includes receptors for siderophores, which were also potentially detected in the samples as well as different biologically important metals. Based on these observations, our hypothesis that the OMVs from *X. citri* can be used for sharing resources in microbial communities is also supported by the observation that the vesicles’ contents can be assimilated and used for microbial growth. At last, another potential resource packaged with the OMVs is their esterase and protease activities, which can release nutrients from the plant host tissue and help to promote microbial colonization, potentially facilitating infection.

This work further establishes the association of OMVs with the acquisition and sharing of nutrient molecules and ions in microbial communities. Microbial interactions can be important driving forces shaping community structure in oligotrophic habitats such as leaf surfaces (Schlechter et al., 2019). The balance between this apparently cooperative behavior with *Xanthomonas*’ notorious competitive proclivities conferred by its bactericidal type IV secretion system (Sgro et al., 2019) may be especially significant for co-occurring epiphytic bacteria and their own particular interactions with the plant (Hassani et al., 2018). Further research on this possible indirect modulation of host physiology could reveal unexplored processes emerging from a pathogen aptly manipulating microbial interaction networks with its diverse suite of secretions systems.

## Materials and Methods

### Bacterial cultures and growth conditions

For all experiments, *Xanthomonas citri* pv. *citri* strain 306 (*X. citri*) was first grown in liquid LB medium (tryptone, 10 g l^-1^; yeast extract, 5 g l^-1^; NaCl, 10 g l^-1^) at 30 °C to OD 0.3 at 600 nm. The cultures were then inoculated on different solid culture media and incubated at 30 °C for 3 days. SB medium (yeast extract, 5 g l^-1^; peptone, 5 g l^-1^; glutamic acid, 1 g l^-1^; sucrose, 5 g l^-1^; pH 7) with 1.5% of agar (w/v) (Ou, 1985), was used for the production and purification of OMVs.

### Purification of OMVs

*X. citri* colonies grown on SB plates at 30 °C for 3 days were scraped from the agar surfaces and suspended in phosphate buffered saline, PBS (NaCl, 8 g l^-1^; KCl, 0.2 g l^-1^; Na_2_HPO_4_, 1.44 g l^-1^; KH_2_PO_4_, 0.24 g l^-1^). After homogenization of the suspension, cells were precipitated by multiple centrifugations steps (10,000 - 30,000 × g at 4 °C, Beckman Avanti J-30I) until the supernatant appeared clean. Then, the samples were ultracentrifuged at 100,000 × g at 4 °C for at least 2 hours to pelletize the OMVs. The pellets were resuspended in a small volume of PBS and filtered through a 0.22 μm syringe filter to remove remaining cells inside a laminar flow hood. The samples were aseptically manipulated from this step on. The filtered OMVs were further purified by being loaded at the bottom of a filtered OptiPrep (Sigma) density gradient (35 to 0% in PBS) and ultracentrifuged at 200,000 × g for at least 12 hours at 4 °C (Beckman Optima XL-100K). The corresponding clear yellow band was collected, diluted in PBS, and pelletized again at 100,000 × g for 2 hours to wash out the density gradient medium. Absence of contamination was determined by lack of growth on LB plates incubated at 30 °C. DLS (Malvern Zetasizer) was used to characterize the size of the recovered OMVs. Total proteins in purified samples were quantified by a Qubit 3.0 fluorometer (Thermo Scientific).

### Sodium dodecyl sulfate-polyacrylamide gel electrophoresis (SDS-PAGE)

Purified OMV samples were added with SDS-PAGE reducing sample buffer and treated at 90 °C for 10 minutes. Proteins were separated in 15% Tris-Glycine SDS-PAGE gels and stained with Coomassie Brilliant Blue.

### Negative stain transmission electron microscopy (TEM)

Samples were applied to glow-discharged carbon film-coated copper grids (400 Mesh, CF400-Cu, Electron Microscopy Sciences), washed with Milli-Q ultrapure water, and negatively stained with uranyl acetate 2% (w/v), blotting on filter paper after each step. A FEI Tecnai G20 200 kV TEM (Department of Cell and Developmental Biology, Institute of Biomedical Sciences, University of São Paulo) or a JEOL JEM 2100 200 kV TEM (Institute of Chemistry, University of São Paulo) were used for image acquisition.

### Liquid chromatography-tandem mass spectrometry lipidomics

Lipids were extracted by the Bligh and Dyer method (Bligh and Dyer, 1959), using ethanol-washed glass tubes and glass Pasteur pipettes for all steps. 100 μl of the samples were added to 400 μl of PBS (50 mM) containing 100 μM of deferoxamine. In the same tubes, 200 μl of a mix of internal standards (Avanti Polar Lipids and Sigma) and 300 μl of butylated hydroxytoluene (BHT) in methanol were added. The samples were then mixed with chloroform/ethyl acetate solution (4:1) and vortexed for 1 minute. Next, the tubes were centrifuged at 1,500 × g for 2 minutes at 4 °C and the organic phase at the bottom was collected and transferred to a clean vial. The solvent was dried under a flow of N_2_ and the lipids were resuspended in 100 μl of isopropyl alcohol. The samples were stored at -80 °C before being analyzed by a previously established untargeted lipidomic method (Chaves-Filho et al., 2019).

### Sample preparation for proteomics analysis

For in-solution digestion, OMV samples were boiled for 10 minutes before the proteins were precipitated with ethanol/acetone, and dissolved in urea 8 M in NH_4_HCO_3_ 100 mM. Dithiothreitol (DTT) was added to a final concentration of 10 mM, and the samples were incubated for 30 min at 37 °C. The samples were cooled down, iodoacetamide was added to a final concentration of 40 mM, and the samples were then incubated for 30 min at room temperature in the dark. DTT was added again, followed by digestion buffer (NH_4_HCO_3_ 50 mM in a solution of 10% acetonitrile - ACN) to dilute ten times the urea concentration. Trypsin was added to digestion buffer for a final trypsin to protein ratio of 1:50, and the solution was incubated overnight at 37 °C. The digestion was stopped by the addition of formic acid (FA).

For in-gel digestion (GeLC approach), the gel bands were completely destained, treated with 10 mM DTT at 56 °C for 45 min, 55 mM IAA at room temperature for 30 min in the dark and digested at 37 °C for 16 hrs with 2 µg sequencing grade modified trypsin, Porcine (Promega). The resultant peptides were extracted in 40% ACN/0.1% TFA into fresh Protein LoBind® microtubes, dried down by vacuum centrifugation, and resuspended in 50 µL 0.1% TFA. Peptide samples obtained from the in-solution and in-gel digestions were desalted using C18 disks packed in a p200 pipette tip. Peptides were eluted with 50% ACN and dried down.

### Nano-flow liquid chromatography-tandem mass spectrometry-based proteomics

Tryptic peptides were resuspended in 0.1% FA and analyzed using an EASY-nLC system (Thermo Scientific) coupled to LTQ-Orbitrap Velos mass spectrometer (Thermo Scientific) at the Core Facility for Scientific Research at the University of São Paulo (CEFAP-USP/BIOMASS). The peptides were loaded onto a C18 PicoFrit column (C18 PepMap, 75 µm id × 10 cm, 3.5 µm particle size, 100 Å pore size; New Objective, Ringoes, NJ, USA) and separated with a gradient from 100% mobile phase A (0.1% FA) to 34% phase B (0.1% FA, 95% ACN) during 60 min, 34%–95% in 15 min, and 5 min at 95% phase B at a constant flow rate of 250 nL/min. The LTQ-Orbitrap Velos was operated in positive ion mode with data-dependent acquisition. The full scan was obtained in the Orbitrap with an automatic gain control target value of 10^6^ ions and a maximum fill time of 500 ms. Each precursor ion scan was acquired at a resolution of 60,000 FWHM in the 400–1500 m/z mass range. Peptide ions were fragmented by CID MS/MS using a normalized collision energy of 35%. The 20 most abundant peptide were selected for MS/MS and dynamically excluded for a duration of 30s. All raw data were accessed in the Xcalibur software (Thermo Scientific).

### Proteomics data analysis

Raw data were processed with MaxQuant (Tyanova et al., 2016) using Andromeda search engine against the SwissProt *Xanthomonas axonopodis* pv. *citri* (strain 306) database (4354 entries downloaded from UniProt.org, Jan/2021) with common contaminants for protein identification. Database searches were performed with the following parameters: precursor mass tolerance of 10 ppm, product ion mass tolerance of 0.6 Da; trypsin cleavage with two missed cleavage allowed; carbamidomethylation of cysteine (57.021 Da) was set as a fixed modification, and oxidation of methionine (15.994 Da) and protein N-terminal acetylation (42.010 Da) were selected as variable modifications. All identifications were filtered to achieve a protein peptide and PSMs, false discovery rate (FDR) of less than 1%, and a minimum of one unique peptide was required for protein identification. Protein quantification was based on the MaxQuant label-free algorithm using both unique and razor peptides for protein quantification. Protein abundance was assessed on label-free protein quantification (LFQ) based on extracted ion chromatogram area of the precursor ions activating the matching between run features. Intensity based absolute quantification (iBAQ) values were used to calculate the relative protein abundance within samples. MS data have been submitted to the PRIDE repository, project accession: PXD025405, username, password: MyMyVfmr.

Statistical enrichment analyses of Pfam and InterPro domains and FDR calculations were obtained from the STRING database (Franceschini et al., 2013). PSORTb 3.0 was used for subcellular localization prediction of the identified proteins (Yu et al., 2010), followed by manual curation based on sequence annotations, and SignalP 5.0 was used for predicting protein secretion mechanisms (Armenteros et al., 2019).

### Elemental analysis by Triple Quadrupole Inductively Coupled Plasma-Mass Spectrometry

Triple Quadrupole Inductively Coupled Plasma-Mass Spectrometry (iCAP TQ ICP-MS, Thermo Fisher Scientific, Bremen, Germany) equipped with a Micro Mist nebulizer (400 µL min^-1^) combined with a cyclonic spray chamber (both obtained from ESI Elemental Service & Instruments GmbH, Mainz, Germany) and an auto-sampler ASX-560 (Teledyne CETAC Technologies, Omaha, NE, USA) was used to perform quantitative analysis of the elements in OMVs samples. The instrument was tuned prior to the elemental analysis to obtain the highest sensitivity. The interface was assembled using a nickel sample cone and a nickel skimmer cone with an insert version for high matrix (3.5 mm).

The TQ ICP-MS was operated with 99.999% Argon (Air Products). Helium and oxygen (99.999%, Linde) were used in the collision/reaction cell of the instrument. A screening (survey scan) was performed on the OMV samples and the PBS buffer (method blank) to identify the main chemical elements contained in the sample, recording the full mass spectrum from 4.6 to 245.0 u.. All measurements were performed in triplicate (n=3) according to selected masses showed in **Table S3**. All data were evaluated with Qtegra ISDS software (Thermo Scientific).

Mono-elemental standard solutions were used for calibration curves. Ca, Mn, Fe, Co, Ni, Cu, Zn, and Ba solutions (PlasmaCAL, SCP Science containing 1000 mg l^-1^ each) were used to calibrate these elements. Mg (1000 mg l^-1^, CertiPUR, Merck), Se (1000 mg l^-1^, Wako Pure Chemical Industries), oxalate standard for carbon quantification (10000 mg l^-1^, TraceCERT, Sigma-Aldrich), and Certified Multielement Ion Chromatography Anion Standard Solution for Bromine and Sulfur quantification (10 mg l^-1^, TraceCERT, Sigma-Aldrich) were also used to calibrate these respectively elements. The OMV samples were diluted to 500 μl with PBS buffer prior to TQ ICP-MS analysis and PBS was used as a method blank. **Table S4** displays the main analytical performance characteristics achieved: linear range, sensitivity, limit of detection (LOD), and coefficient of determination.

Instrumental precision was checked by stability tests throughout the analysis (obtaining a relative standard deviation of less than 3% for all analytes) and the accuracy was checked by spike and recovery tests at four different levels of concentration, obtaining acceptable values ranging from 93 to 105%.

### Siderophore detection and bacterial growth assays

The presence of siderophores in the purified OMVs was tested on chrome azurol S (CAS) agar plates (Schwyn and Neilands, 1987), prepared according to Louden et al. (2011). Bacterial growth using purified OMVs as sole carbon sources was assayed in M9 minimal medium without glucose (Na_2_HPO_4_, 6.8 g l^-1^; KH_2_PO_4_, 3 g l^-1^; NH_4_Cl, 1 g l^-1^; NaCl, 0.5 g l^-1^; MgSO_4_, 2 mM; CaCl_2_, 2 mM). About 10^3^ stationary phase cells l^-1^, equivalent to around 10 colony forming units (CFU) for each 10 µl droplet plated, were used as the initial population for the experiments. To the samples, 0, 125, or 375 µg ml^-1^ of total OMV proteins were added, and the tubes were incubated at 30 °C in a thermomixer for 48 h. Aliquots were taken at regular intervals and plated in LB medium for CFU quantification.

### Esterase activity assays

Esterase qualitative assays were performed on either LB plates prepared with 0.5% tributyrin emulsified by sonication (SONICS Vibra-Cell), or NYG plates (peptone, 5 g l^-1^; yeast extract, 5 g l^-1^; glycerol, 20 g l^-1^, agar, 1% (Turner et al., 1984)) containing 1% of Tween 20 and 4 mM of CaCl_2_ (Ramnath et al., 2017). Esterase enzymatic activity was measured colorimetrically with a reaction mixture (100 mM Tris-HCl pH 7.5, 50 mM NaCl) containing 500 µM of *p*NP-C4 or 200 µM of *p*NP-C8 with the addition of 10, 20, or 50 µg ml^-1^ of total proteins of purified OMVs in microplate wells. The reactions were incubated at 30 °C and their absorbance at 400 nm was measured with a SpectraMax Paradigm microplate reader (Molecular Devices) at regular intervals of time during 4 h.

### Protease activity assays

Protease assays were performed with a protease fluorescent detection kit using casein labeled with fluorescein isothiocyanate (FITC) as the substrate following the manufacturer’s instructions (PF0100, Sigma-Aldrich). Briefly, 10 µl of the test samples were added to 40 µl of FITC-casein in incubation buffer and incubated at 30 °C for 6 hours. PBS was used as a blank, and the reactions contained 100, 200, or 300 µg ml^-1^ of total proteins of purified OMVs. For some assays, EDTA-free Pierce Protease Inhibitor (A32965, Thermo Scientific) was added to 300 µg ml^-1^ samples to a final concentration equivalent to the manufacturer’s recommendations (1 tablet for 50 ml of solution). After incubation, undigested substrate was precipitated with the addition of 150 µl of trichloroacetic acid 0.6 N for 30 min at 37 °C. Aliquots of the supernatants containing FITC-labeled fragments were diluted in assay buffer and analyzed in a black 96-well microplate with a SpectraMax Paradigm microplate reader (Molecular Devices). Relative fluorescence units (RFU) were measured with excitation at 485 nm and detection at 535 nm. All samples presented RFU measurements substantially above 120% of the value obtained with the blank (data not shown), which is considered significant according to the kit’s manufacturer.

## Acknowledgements

The authors would like to thank Roberto Cabado Modia Junior and Alfredo Duarte for the technical assistance at the electron microscopy facilities, Thais Viggiani Santana for the assistance at the proteomics facility, and Tania Geraldine Churasacari Vinces for the assistance with the esterase activity experiments.

We thank the Core Facility for Scientific Research – University of São Paulo (CEFAP-USP/BIOMASS) for the proteomic analysis. The authors acknowledge financial support from the São Paulo Research Foundation (FAPESP): grants 2019/00195-2 and 2020/04680-0 to CRG, 2017/17303-7 to CSF, 2014/06863-3, 2018/18257-1 and 2018/15549-1 to GP, 2013/07937-8 to SM, 2016/09047-8 to RFdS, 2017/20752-8 to the EMU TQ ICP-MS facility at IPT, and scholarships 2018/21076-9 to GGA, 2017/24301-0 to MMC, 2017/10611-8 to ADP, 2019/12234-2 to EEL, and 2017/13804-1 to AI. The authors also acknowledge financial support from the Coordenação de Aperfeiçoamento de Pessoal de Nível Superior (CAPES) in the form of scholarships to MMY, IT, and GGA (88887.336498/2019-00), and the Brazilian National Council for Scientific and Technological Development (CNPq) for RP grants 380490/2018-8 and 380939/2020-7.

---

**Data Set S1**. Proteomic data for purified *X. citri* OMVs, containing details of the filtered proteins identified for the duplicate of in-solution digestions, including their iBAQ values (XLSX file).

**Data Set S2**. Proteomic data for purified *X. citri* OMVs, containing details of the filtered proteins identified for the in-gel digestion, including their iBAQ values (XLSX file).

**Table S1.** Combination of the top 100 most abundant proteins determined by their iBAQ values from the different purified *X. citri* OMV samples (gel bands and a replicate of samples in solution, **Fig. 4**), resulting in a list of 163 non-redundant proteins. UniProt annotations are presented for each sequence.

| UniProt ID | Gene names | Locus tags | Protein names (UniProt) | Pfam domains | InterPro domains |
| --- | --- | --- | --- | --- | --- |
| Q8PRF7 |  | XAC0006 | Peptidase_M48 domain-containing protein | PF01435 | IPR001915 |
| Q8PRF6 |  | XAC0007 | TPR_REGION domain-containing protein | PF13181 | IPR013026; IPR011990; IPR019734 |
| Q8PRF4 | exbB | XAC0009 | Biopolymer transport ExbB protein | PF01618 | IPR002898 |
| Q8PRE0 | ctp | XAC0023 | Carboxyl-terminal protease | PF13180; PF03572 | IPR029045; IPR001478; IPR036034; IPR004447; IPR005151 |
| Q8PRD3 | egl | XAC0030 | Cellulase | PF00150 | IPR001547; IPR018087; IPR017853 |
| Q8PRC7 |  | XAC0036 | Uncharacterized protein |  |  |
| Q8PR40 | yncD | XAC0126 | Iron transporter | PF07715; PF00593 | IPR039426; IPR012910; IPR037066; IPR000531; IPR036942 |
| Q8PQZ3 | fpvA | XAC0176 | Ferripyoverdine receptor | PF07715; PF00593 | IPR012910; IPR037066; IPR039423; IPR000531; IPR036942; IPR010105 |
| Q8PQZ2 |  | XAC0177 | PNPLA domain-containing protein | PF01734 | IPR016035; IPR002641 |
| Q8PQX9 |  | XAC0190 | Uncharacterized protein |  | IPR011990 |
| Q8PQW2 | yojM | XAC0209 | Superoxide dismutase [Cu-Zn] (EC 1.15.1.1) | PF00080 | IPR036423; IPR024134; IPR018152; IPR001424 |
| Q8PQU8 |  | XAC0223 | Uncharacterized protein |  | IPR026364; IPR023614 |
| Q8PQT9 |  | XAC0232 | Uncharacterized protein | PF13698 | IPR025294 |
| Q8PQQ0 |  | XAC0272 | Uncharacterized protein |  |  |
| Q8PQN4 |  | XAC0289 | Uncharacterized protein |  | IPR016980; IPR029063 |
| Q8NL21 | rplM | XAC0487 | 50S ribosomal protein L13 | PF00572 | IPR005822; IPR005823; IPR023563; IPR036899 |
| Q8PPZ1 | groL | XAC0542 | 60 kDa chaperonin (GroEL protein) (Protein Cpn60) | PF00118 | IPR018370; IPR001844; IPR002423; IPR027409; IPR027413; IPR027410 |
| Q8PPR2 |  | XAC0623 | Uncharacterized protein | PF04338 | IPR007433 |
| Q8PPM3 | rlpA | XAC0663 | Endolytic peptidoglycan transglycosylase RlpA (EC 4.2.2.-) | PF03330; PF05036 | IPR034718; IPR009009; IPR036908; IPR012997; IPR007730; IPR036680 |
| Q8PPM2 | dacC | XAC0664 | Serine-type D-Ala-D-Ala carboxypeptidase (EC 3.4.16.4) | PF07943; PF00768 | IPR012338; IPR015956; IPR018044; IPR012907; |
|  |  |  |  |  | IPR037167; IPR001967 |
| **Q8PPK9** |  | XAC0677 | Uncharacterized protein | PF10001 | IPR018718 |
| **Q8PPK4** |  | XAC0682 | BON domain-containing protein | PF04972 | IPR007055; IPR014004 |
| **Q8PPJ6** | fecA | XAC0690 | TonB-dependent receptor | PF07715;<br>PF00593 | IPR012910; IPR037066;<br>IPR039423; IPR000531;<br>IPR036942 |
| **Q8PPH0** | fyuA | XAC0716 | TonB-dependent receptor | PF07715;<br>PF00593 | IPR012910; IPR037066;<br>IPR000531; IPR036942;<br>IPR010104 |
| **Q8PPD9** |  | XAC0747 | Uncharacterized protein |  | IPR023614 |
| **Q8PPC1** |  | XAC0765 | Uncharacterized protein | PF04348 | IPR007443; IPR028082 |
| **Q8PP23** | surA | XAC0865 | Chaperone SurA (Peptidyl-prolyl cis-trans isomerase SurA) (PPIase SurA) (EC 5.2.1.8) (Rotamase SurA) | PF00639;<br>PF09312 | IPR000297; IPR023034;<br>IPR015391; IPR027304 |
| **Q8PP00** | gfo | XAC0888 | Glucose-fructose oxidoreductase | PF01408;<br>PF02894 | IPR004104; IPR008354;<br>IPR036291; IPR000683 |
| **Q8PNT4** | rplK | XAC0961 | 50S ribosomal protein L11 | PF00298;<br>PF03946 | IPR000911; IPR036796;<br>IPR006519; IPR020783;<br>IPR036769; IPR020785;<br>IPR020784 |
| **Q8PNS2** | rplW | XAC0974 | 50S ribosomal protein L23 | PF00276 | IPR012677; IPR012678;<br>IPR013025 |
| **Q8PNS1** | rplB | XAC0975 | 50S ribosomal protein L2 | PF00181;<br>PF03947 | IPR012340; IPR022666;<br>IPR014722; IPR002171;<br>IPR005880; IPR022669;<br>IPR022671; IPR014726;<br>IPR008991 |
| **Q8NKY0** | rplV | XAC0977 | 50S ribosomal protein L22 | PF00237 | IPR001063; IPR018260;<br>IPR036394; IPR005727 |
| **Q8PNR8** | rplP | XAC0979 | 50S ribosomal protein L16 | PF00252 | IPR016180; IPR036920;<br>IPR000114; IPR020798 |
| **Q8NL02** | rplN | XAC0982 | 50S ribosomal protein L14 | PF00238 | IPR036853; IPR000218;<br>IPR005745; IPR019972 |
| **Q8PNR4** | rplE | XAC0984 | 50S ribosomal protein L5 | PF00281;<br>PF00673 | IPR002132; IPR020930;<br>IPR031309; IPR020929;<br>IPR022803; IPR031310 |
| **Q8PNR3** | rpsH | XAC0986 | 30S ribosomal protein S8 | PF00410 | IPR000630; IPR035987 |
| **Q8PNR1** | rplR | XAC0988 | 50S ribosomal protein L18 | PF00861 | IPR005484; IPR004389 |
| **Q8PNR0** | rpmD | XAC0990 | 50S ribosomal protein L30 | PF00327 | IPR036919; IPR005996;<br>IPR016082 |
| **Q8PNQ9** | rplO | XAC0991 | 50S ribosomal protein L15 | PF00828 | IPR036227; IPR030878;<br>IPR005749; IPR001196;<br>IPR021131 |
| **Q8NKX3** | rpsM | XAC0993 | 30S ribosomal protein S13 | PF00416 | IPR027437; IPR001892;<br>IPR010979; IPR019980;<br>IPR018269 |
| **P0A0Y0** | rpsD | XAC0995 | 30S ribosomal protein S4 | PF00163;<br>PF01479 | IPR022801; IPR001912;<br>IPR005709; IPR018079;<br>IPR002942; IPR036986 |
| **Q8PNQ7** | rplQ | XAC0997 | 50S ribosomal protein L17 | PF01196 | IPR000456; IPR036373 |
| **Q8PNP2** | mopB | XAC1012 | Outer membrane protein | PF13505;<br>PF00691 | IPR011250; IPR027385;<br>IPR006664; IPR006665;<br>IPR036737; IPR028974 |
| **Q8PNJ7** |  | XAC1062 | Uncharacterized protein |  |  |
| **Q8PNF8** | slp | XAC1113 | Outer membrane protein Slp | PF03843 | IPR004658 |
| **Q8PND0** | fyuA | XAC1143 | TonB-dependent receptor | PF07715;<br>PF00593 | IPR012910; IPR039423;<br>IPR000531; IPR036942 |
| **Q8PNB0** |  | XAC1163 | Uncharacterized protein |  |  |
| **Q8PN49** | minE | XAC1224 | Cell division topological specificity factor | PF03776 | IPR005527; IPR036707 |
| **Q8PN43** |  | XAC1230 | Uncharacterized protein |  | IPR011256 |
| **Q8PN35** |  | XAC1238 | Endo/exonuclease/phosphatase domain-containing protein | PF03372 | IPR036691; IPR005135 |
| **Q8PN33** |  | XAC1240 | Uncharacterized protein | PF13202;<br>PF13499 | IPR011992; IPR018247;<br>IPR002048 |
| **Q8PN25** | rplU | XAC1248 | 50S ribosomal protein L21 | PF00829 | IPR036164; IPR028909;<br>IPR001787; IPR018258 |
| **Q8PMV4** | mucD | XAC1321 | Periplasmic serine endoprotease DegP-like (EC 3.4.21.107) | PF13180 | IPR001478; IPR036034;<br>IPR011782; IPR009003;<br>IPR001940 |
| **Q8PMV1** |  | XAC1324 | Uncharacterized protein | PF16137 | IPR032314 |
| **Q8PMS7** |  | XAC1349 | Serine protease | PF03797;<br>PF12951;<br>PF00082 | IPR005546; IPR036709;<br>IPR013425; IPR000209;<br>IPR036852; IPR023827;<br>IPR023828; IPR015500;<br>IPR034061 |
| **Q8PMP7** |  | XAC1379 | Uncharacterized protein | PF10099 | IPR018764 |
| **Q8PMM9** |  | XAC1397 | Alginate_exp domain-containing protein | PF13372 | IPR025388 |
| **Q8PML3** | oma | XAC1413 | Outer membrane protein assembly factor BamA | PF01103;<br>PF07244 | IPR000184; IPR010827;<br>IPR039910; IPR023707;<br>IPR034746 |
| **Q8PMJ3** |  | XAC1434 | CASH domain-containing protein | PF13229 | IPR039448; IPR006633;<br>IPR022441; IPR006626;<br>IPR012334; IPR011050 |
| **Q8PMJ2** | fhuA | XAC1435 | Iron receptor | PF07715;<br>PF00593 | IPR012910; IPR037066;<br>IPR039423; IPR000531;<br>IPR036942; IPR010917;<br>IPR010105 |
| **Q8PMH2** | dcp | XAC1456 | Peptidyl-dipeptidase | PF01432 | IPR034005; IPR024077;<br>IPR001567 |
| **Q8PMG5** |  | XAC1463 | Phospholipase A1 (EC 3.1.1.32) (EC | PF02253 | IPR003187; IPR036541 |
|  |  |  | 3.1.1.4) (Phosphatidylcholine 1-acylhydrolase) |  |  |
| **Q8PMG3** | pcp | XAC1466 | Peptidoglycan-associated outer membrane lipoprotein | PF05433 | IPR008816 |
| **Q8PMF0** |  | XAC1479 | OmpA family protein | PF13488;<br>PF00691 | IPR039567; IPR006664;<br>IPR006665; IPR006690;<br>IPR036737 |
| **Q8PMD2** |  | XAC1497 | Uncharacterized protein |  |  |
| **Q8PMB6** | smpA | XAC1516 | Outer membrane protein assembly factor BamE | PF04355 | IPR026592; IPR037873;<br>IPR007450 |
| **Q8PM83** | btuE | XAC1549 | Glutathione peroxidase | PF00255 | IPR000889; IPR029759;<br>IPR036249 |
| **Q8PM82** | fkpA | XAC1550 | Peptidyl-prolyl cis-trans isomerase (EC 5.2.1.8) | PF00254;<br>PF01346 | IPR001179; IPR000774;<br>IPR036944 |
| **Q8PM54** | oprO | XAC1579 | Polyphosphate-selective porin O | PF07396 | IPR023614; IPR010870 |
| **Q8NL26** |  | XAC1585 | Peptidyl-prolyl cis-trans isomerase (EC 5.2.1.8) | PF00254;<br>PF01346 | IPR001179; IPR000774;<br>IPR036944 |
| **Q8PM13** | rpsR | XAC1621 | 30S ribosomal protein S18 | PF01084 | IPR001648; IPR018275;<br>IPR036870 |
| **Q8PLS7** |  | XAC1712 | DUF218 domain-containing protein | PF02698 | IPR003848; IPR014729 |
| **Q8PLR1** | nlpD | XAC1728 | Lipoprotein | PF01476;<br>PF01551 | IPR011055; IPR018392;<br>IPR036779; IPR016047 |
| **Q8PLN4** |  | XAC1761 | Uncharacterized protein |  |  |
| **Q8PL93** | cirA | XAC1910 | TonB-dependent receptor | PF07715;<br>PF00593 | IPR012910; IPR037066;<br>IPR000531; IPR010104 |
| **Q8PKZ8** | lolA | XAC2008 | Outer-membrane lipoprotein carrier protein | PF03548 | IPR029046; IPR004564;<br>IPR018323 |
| **Q8PKZ0** | pilF | XAC2017 | Fimbrial biogenesis protein |  | IPR013360; IPR013026;<br>IPR011990; IPR019734 |
| **Q8PKY7** | bamB | XAC2020 | Outer membrane protein assembly factor BamB | PF13360 | IPR017687; IPR018391;<br>IPR002372; IPR011047;<br>IPR015943 |
| **Q8PK64** |  | XAC2312 | TonB_dep_Rec domain-containing protein | PF00593 | IPR039426; IPR013784;<br>IPR000531 |
| **Q8PK57** |  | XAC2319 | Uncharacterized protein |  |  |
| **Q8PK24** |  | XAC2353 | Uncharacterized protein |  |  |
| **Q8PJM6** | rpfN | XAC2504 | Porin | PF04966 | IPR007049; IPR038673 |
| **Q8PJK8** | ggt | XAC2523 | Gamma-glutamyltranspeptidase |  | IPR043138; IPR000101;<br>IPR043137; IPR029055 |
| **Q8PJK6** |  | XAC2525 | Uncharacterized protein |  |  |
| **Q8PJK0** | btuB | XAC2531 | TonB-dependent receptor | PF07715;<br>PF00593 | IPR012910; IPR037066;<br>IPR000531; IPR010916 |
| **Q8PJH0** |  | XAC2562 | Uncharacterized protein |  |  |
| **Q8PJE3** | rplT | XAC2591 | 50S ribosomal protein L20 | PF00453 | IPR005813; IPR035566 |
| **Q8PJD5** | btuB | XAC2600 | TonB-dependent receptor | PF07715 | IPR012910; IPR037066;<br>IPR010104; IPR010917 |
| **Q8PJC4** |  | XAC2611 | DUF4189 domain-containing protein | PF13827 | IPR025240 |
| **Q8PJB5** | virB9 | XAC2620 | VirB9 protein | PF03524 | IPR010258; IPR033645; IPR038161 |
| **Q8PJ70** | oar | XAC2672 | Oar protein |  | IPR039426; IPR008969; IPR036942 |
| **Q8PJ58** | rpsO | XAC2684 | 30S ribosomal protein S15 | PF00312 | IPR000589; IPR005290; IPR009068 |
| **Q8PJ03** | btuB | XAC2742 | TonB-dependent receptor | PF07715; PF00593 | IPR012910; IPR037066; IPR000531; IPR010917 |
| **Q8PJ02** | oar | XAC2743 | Oar protein | PF07715; PF00593 | IPR039426; IPR013784; IPR012910; IPR037066; IPR000531 |
| **Q8PIY6** | phoA | XAC2759 | Alkaline phosphatase | PF00245 | IPR001952; IPR017850 |
| **Q8PIX3** | bp26 | XAC2772 | Outer membrane protein | PF04402 | IPR007497 |
| **Q8PIX2** | oar | XAC2773 | Oar protein | PF07715; PF00593 | IPR039426; IPR013784; IPR012910; IPR037066; IPR000531 |
| **Q8PIW5** | rlpB | XAC2780 | LPS-assembly lipoprotein LptE | PF04390 | IPR007485 |
| **Q8PIU4** |  | XAC2801 | Uncharacterized protein | PF06629 | IPR010583 |
| **Q8PIR6** | phuR | XAC2829 | Outer membrane hemin receptor | PF07715; PF00593 | IPR039426; IPR012910; IPR037066; IPR000531; IPR036942 |
| **Q8PIF7** | fhuA | XAC2941 | TonB-dependent receptor | PF07715; PF00593 | IPR012910; IPR037066; IPR039423; IPR000531; IPR036942; IPR010105 |
| **Q8PIF2** |  | XAC2946 | Uncharacterized protein | PF10670 | IPR019613 |
| **Q8PIE7** | comEA | XAC2951 | DNA transport competence protein |  | IPR004509; IPR010994 |
| **Q8PIE0** |  | XAC2958 | Uncharacterized protein | PF09839 | IPR018642 |
| **Q8PID5** |  | XAC2963 | Uncharacterized protein | PF11306 | IPR021457 |
| **Q8PI48** | btuB | XAC3050 | TonB-dependent receptor | PF07715; PF00593 | IPR012910; IPR037066; IPR000531; IPR036942 |
| **Q8PI41** | bla | XAC3057 | Beta-lactamase | PF00144 | IPR001466; IPR012338 |
| **Q8PI27** | iroN | XAC3071 | TonB-dependent receptor | PF07715; PF00593 | IPR012910; IPR037066; IPR000531; IPR010104 |
| **Q8PHZ0** |  | XAC3108 | Uncharacterized protein |  | IPR011990 |
| **Q8PHV9** | cpoB | XAC3140 | Cell division coordinator CpoB | PF16331; PF13525 | IPR039565; IPR034706; IPR014162; IPR013026; IPR011990; IPR019734; IPR032519 |
| **Q8PHV8** | ompP6 | XAC3141 | Peptidoglycan-associated protein | PF00691 | IPR006664; IPR006665; IPR036737; IPR039001; IPR014169 |
| **Q8PHV7** | tolB | XAC3142 | Tol-Pal system protein TolB | PF07676; PF04052 | IPR011042; IPR011659; IPR014167; IPR007195; IPR036752 |
| **Q8PHV5** | tolR | XAC3144 | Tol-Pal system protein TolR | PF02472 | IPR003400; IPR014168 |
| **Q8PHV4** | tolQ | XAC3145 | Tol-Pal system protein TolQ | PF01618 | IPR002898; IPR014163 |
| **Q8PHU4** |  | XAC3155 | Uncharacterized protein | PF11218 | IPR021381 |
| **Q8PHT7** | bla | XAC3162 | Beta-lactamase (EC 3.5.2.6) |  | IPR012338; IPR000871; IPR023650; IPR006311 |
| **Q8PHT1** | bfeA | XAC3168 | Ferric enterobactin receptor | PF07715; PF00593 | IPR012910; IPR037066; IPR000531; IPR010916; IPR036942 |
| **Q8PHT0** | bfeA | XAC3169 | Ferric enterobactin receptor | PF07715; PF00593 | IPR012910; IPR037066; IPR000531; IPR036942 |
| **Q8PHS3** | fecA | XAC3176 | Citrate-dependent iron transporter | PF07715; PF00593 | IPR012910; IPR037066; IPR000531; IPR036942; IPR010105 |
| **Q8PHQ5** | btuB | XAC3194 | Outer membrane receptor for transport of vitamin B | PF07715; PF00593 | IPR010101; IPR039426; IPR012910; IPR037066; IPR000531; IPR036942 |
| **Q8PHP1** | bfeA | XAC3207 | Ferric enterobactin receptor | PF07715; PF00593 | IPR012910; IPR037066; IPR000531; IPR010916; IPR036942 |
| **Q8PHN1** | comL | XAC3218 | Outer membrane protein assembly factor BamD | PF13525 | IPR017689; IPR039565; IPR013026; IPR011990 |
| **Q8PHL0** | fimA | XAC3241 | Fimbrillin | PF07963; PF00114 | IPR012902; IPR001082 |
| **Q8PHF7** | estA | XAC3300 | Lipase | PF03797; PF00657 | IPR005546; IPR036709; IPR001087; IPR017186; IPR036514 |
| **Q8PHE6** | iroN | XAC3311 | TonB-dependent receptor | PF07715; PF00593 | IPR012910; IPR037066; IPR006311; IPR000531; IPR036942; IPR010104 |
| **Q8PHC5** | fecA | XAC3334 | TonB-dependent receptor | PF07715; PF00593 | IPR039426; IPR012910; IPR037066; IPR000531; IPR036942 |
| **Q8PHA8** |  | XAC3351 | Uncharacterized protein |  |  |
| **Q8PHA5** | ompW | XAC3354 | Outer membrane protein W | PF03922 | IPR011250; IPR005618 |
| **Q8PHA4** | omp21 | XAC3355 | Outer membrane protein | PF03922 | IPR011250; IPR005618 |
| **Q8PH89** | fhuE | XAC3370 | Outer membrane receptor for ferric iron uptake | PF07715; PF00593 | IPR012910; IPR037066; IPR039423; IPR000531; IPR036942; IPR010105 |
| **Q8PH16** | btuB | XAC3444 | TonB-dependent receptor | PF07715; PF00593 | IPR012910; IPR037066; IPR000531; IPR036942 |
| **Q8PGZ9** | tolC | XAC3463 | TolC protein | PF02321 | IPR003423; IPR010130 |
| **Q8PGZ0** | oprO | XAC3472 | Polyphosphate-selective porin O | PF07396 | IPR023614; IPR010870 |
| **Q8PGX3** | fyuA | XAC3489 | TonB-dependent receptor | PF07715; PF00593 | IPR012910; IPR037066; IPR039423; IPR000531; IPR036942 |
| **Q8PGW4** | fhuE | XAC3498 | Outer membrane receptor for ferric iron uptake | PF07715; PF00593 | IPR012910; IPR037066; IPR039423; IPR000531; |
|  |  |  |  |  | IPR036942; IPR010917;<br>IPR010105 |
| **Q8PGU1** |  | XAC3525 | Uncharacterized protein |  |  |
| **Q8PGL1** | uptE | XAC3605 | Outer membrane protein |  | IPR036737 |
| **Q8PGL0** | uptD | XAC3606 | Outer membran protein | PF14346 | IPR025511 |
| **Q8PGJ6** | pfeA | XAC3620 | Siderophore receptor protein | PF07715;<br>PF00593 | IPR039426; IPR012910;<br>IPR037066; IPR000531;<br>IPR036942; IPR010917;<br>IPR010105 |
| **Q8PGG6** | atpG | XAC3650 | ATP synthase gamma chain (ATP synthase F1 sector gamma subunit) | PF00231 | IPR035968; IPR000131;<br>IPR023632 |
| **Q8PGG0** |  | XAC3657 | Uncharacterized protein |  | IPR011250 |
| **Q8PGF7** |  | XAC3660 | Uncharacterized protein |  |  |
| **Q8PGF3** | ompW | XAC3664 | Outer membrane protein | PF03922 | IPR011250; IPR005618 |
| **Q8PGF0** |  | XAC3667 | Lipoprotein | PF03180 | IPR004872 |
| **Q8PGC9** | dadA | XAC3688 | D-amino acid dehydrogenase (EC 1.4.99.-) | PF01266 | IPR023080; IPR006076;<br>IPR036188 |
| **Q8PFX5** | amaA | XAC3847 | N-acyl-L-amino acid amidohydrolase | PF07687;<br>PF01546 | IPR017439; IPR036264;<br>IPR002933; IPR011650 |
| **Q8PFW2** |  | XAC3860 | N-acetylmuramoyl-L-alanine amidase | PF01510 | IPR036505; IPR002502 |
| **Q8PFV4** | yliI | XAC3868 | Dehydrogenase | PF07995 | IPR011042; IPR012938;<br>IPR011041 |
| **P66535** | rpsU | XAC3872 | 30S ribosomal protein S21 | PF01165 | IPR001911; IPR018278;<br>IPR038380 |
| **Q8PFR1** |  | XAC3917 | SPOR domain-containing protein | PF05036 | IPR007730; IPR036680 |
| **Q8PFQ2** |  | XAC3926 | OMP_b-brl domain-containing protein | PF13505 | IPR011250; IPR027385 |
| **Q8PFK1** | htrA | XAC3980 | Periplasmic serine endoprotease DegP-like (EC 3.4.21.107) | PF13180 | IPR001478; IPR036034;<br>IPR011782; IPR009003;<br>IPR001940 |
| **Q8PFK0** |  | XAC3981 | Uncharacterized protein |  |  |
| **Q8PFH3** | ecnA | XAC4008 | Entericidin A | PF08085 | IPR012556 |
| **Q8PFD5** | iroN | XAC4048 | TonB-dependent receptor | PF07715;<br>PF00593 | IPR012910; IPR037066;<br>IPR000531; IPR010104 |
| **P66160** | rpmB | XAC4159 | 50S ribosomal protein L28 | PF00830 | IPR034704; IPR026569;<br>IPR037147; IPR001383 |
| **Q8PEX1** |  | XAC4219 | Ysc84 domain-containing protein | PF04366 | IPR007461 |
| **Q8PER7** |  | XAC4273 | OmpA-related protein | PF00593 | IPR039426; IPR013784;<br>IPR000531 |
| **Q8PER6** |  | XAC4274 | OmpA-related protein | PF00593 | IPR039426; IPR013784;<br>IPR000531 |
| **Q8PEK7** | yrbC | XAC4342 | Toluene tolerance protein | PF05494 | IPR008869; IPR042245 |
| **Q8PEK5** | vacJ | XAC4344 | Lipoprotein | PF04333 | IPR007428 |
| **Q8PRJ3** | virB9 | XACb0039 | VirB9 protein | PF03524 | IPR010258; IPR014148;<br>IPR033645; IPR038161 |
| **Q8NL05** | rpsN |  | 30S ribosomal protein S14 | PF00253 | IPR001209; IPR043140;<br>IPR023036 |

**Table S2.**
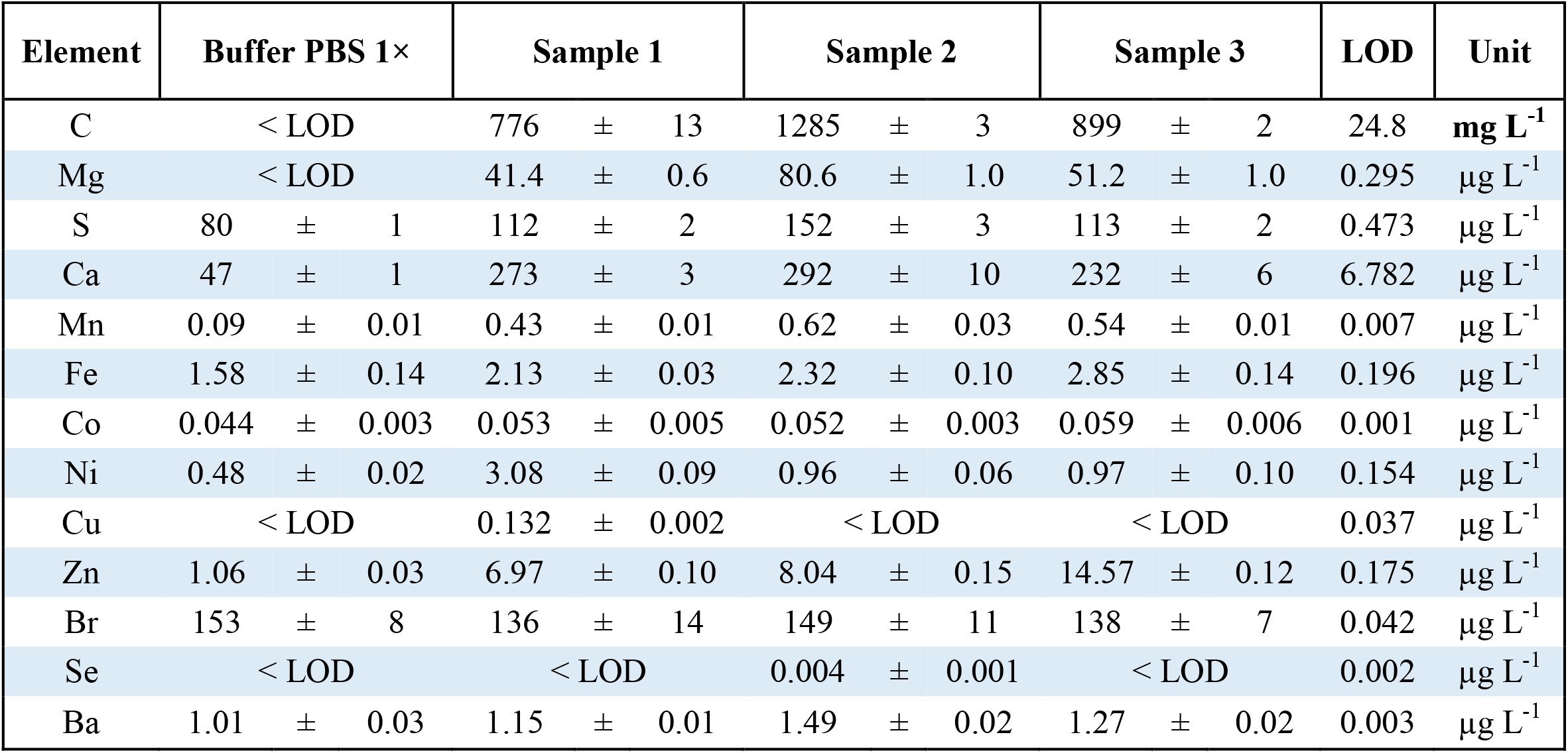
Results from the TQ ICP-MS elemental analysis of samples containing purified OMV suspended in PBS. The data for **Fig. 6B** were obtained by subtracting the background concentration of each element in PBS and normalizing the values for each sample based on their respective carbon content. See also **Table S3** and **Table S4** for experimental details. LOD: limit of detection.

**Table S3.** Mass values defined in the quadrupoles for the TQ ICP-MS elemental analysis.

| Element | Isotope | Analysis Mode | Q<sub>1</sub> mass | Q<sub>2</sub> filled with | Q<sub>3</sub> mass | LOD | Unit | Linear Range | Sensitivity (cps L µg<sup>-1</sup>) | R<sup>2</sup> |
| --- | --- | --- | --- | --- | --- | --- | --- | --- | --- | --- |
| C | 12 | SQ - KED | --- | Helium | 12 | 24.8 | mg L<sup>-1</sup> | 50 - 1500 | 3.0 × 10<sup>2</sup> | 0.9999 |
| Mg | 24 | SQ - KED | --- | Helium | 24 | 0.295 | µg L<sup>-1</sup> | 25 - 100 | 6.3 × 10<sup>2</sup> | 0.9924 |
| S | 32 | TQ - O2 | 32 | Oxygen | <sup>48</sup>(<sup>32</sup>S.<sup>16</sup>O<sup>+</sup>) | 0.473 | µg L<sup>-1</sup> | 3 - 160 | 1.9 × 10<sup>3</sup> | 0.9987 |
| Ca | 44 | SQ - KED | --- | Helium | 44 | 6.782 | µg L<sup>-1</sup> | 25 - 500 | 4.5 × 10<sup>1</sup> | 0.9997 |
| Mn | 55 | SQ - KED | --- | Helium | 55 | 0.007 | µg L<sup>-1</sup> | 0.01 - 5 | 9.2 × 10<sup>3</sup> | 0.9999 |
| Fe | 57 | SQ - KED | --- | Helium | 57 | 0.196 | µg L<sup>-1</sup> | 0.25 - 5 | 4.1 × 10<sup>2</sup> | 0.9977 |
| Co | 59 | SQ - KED | --- | Helium | 59 | 0.001 | µg L<sup>-1</sup> | 0.01 - 1 | 3.8 × 10<sup>4</sup> | 0.9993 |
| Ni | 60 | SQ - KED | --- | Helium | 60 | 0.154 | µg L<sup>-1</sup> | 0.25 - 10 | 1.2 × 10<sup>4</sup> | 0.9939 |
| Cu | 63 | SQ - KED | --- | Helium | 63 | 0.037 | µg L<sup>-1</sup> | 0.05 - 10 | 3.1 × 10<sup>4</sup> | 0.9999 |
| Zn | 66 | SQ - KED | --- | Helium | 66 | 0.175 | µg L<sup>-1</sup> | 0.25 - 25 | 4.1 × 10<sup>3</sup> | 0.9997 |
| Br | 79 | TQ - O2 | 79 | Oxygen | <sup>95</sup>(<sup>79</sup>Br.<sup>16</sup>O<sup>+</sup>) | 0.042 | µg L<sup>-1</sup> | 50 - 500 | 3.6 × 10<sup>2</sup> | 0.9999 |
| Se | 80 | TQ - O2 | 80 | Oxygen | <sup>96</sup>(<sup>80</sup>Se.<sup>16</sup>O<sup>+</sup>) | 0.002 | µg L<sup>-1</sup> | 0.005 - 1 | 2.1 × 10<sup>3</sup> | 0.9993 |
| Ba | 138 | TQ - O2 | 138 | Oxygen | <sup>154</sup>(<sup>138</sup>Ba.<sup>16</sup>O<sup>+</sup>) | 0.003 | µg L<sup>-1</sup> | 0.05 - 5 | 5.2 × 10<sup>4</sup> | 0.9999 |

**Table S4.** TQ ICP-MS operating conditions.

|  |  |
| --- | --- |
| **RF Power (W)** | **1550** |
| **Argon coolant gas flow (L min<sup>-1</sup>)** | 14 |
| **Argon auxiliary gas flow (L min<sup>-1</sup>)** | 0.8 |
| **Argon nebulizer flow (L min<sup>-1</sup>)** | 1.04 |
| **He flow gas (mL min<sup>-1</sup>)** | 6.57 |
| **O<sub>2</sub> flow gas (mL min<sup>-1</sup>)** | 0.6 |
| **Nebulizer** | MicroMist U-Series 0.4 mL min<sup>-1</sup> |
| **Spray Chamber** | Glass Cyclonic spray chamber |
| **Peristaltic Pump (rpm)** | 40 |
| **Spray chamber temperature (°C)** | 2.7 |
| **Dwell time (s)** | 0.1 |
| **Number of Sweeps** | 10 |

**Fig. S1.**
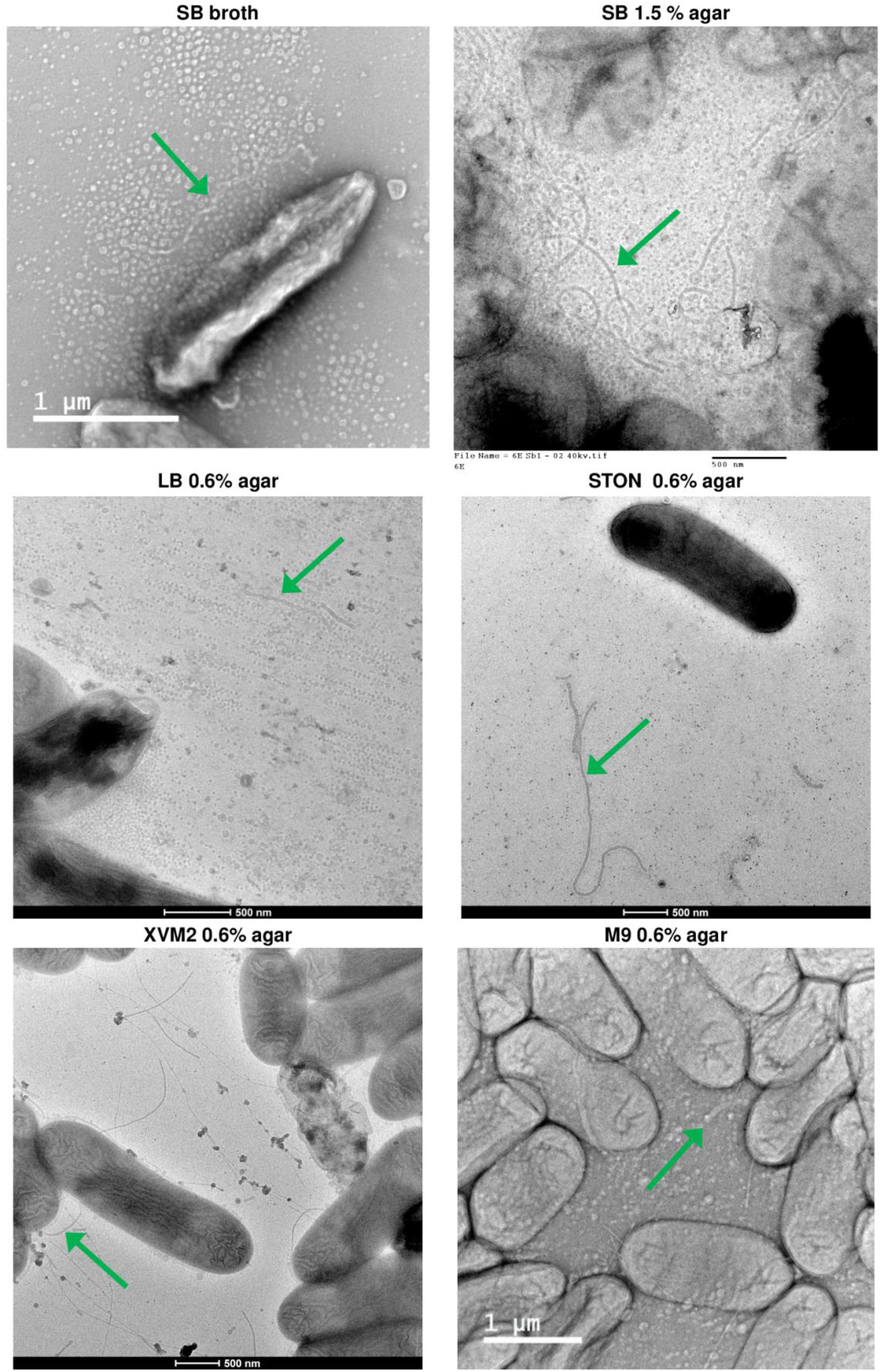
Formation of outer membrane tubes by *X. citri* cells in different culture conditions and media. The tested media include liquid SB, in which the samples were concentrated by ultracentrifugation before being applied to the TEM grids, SB with 1.5% agar (a higher concentration than the 0.6% used for Fig. 1), LB with 0.6% agar, STON with 0.6% agar (Guzzo et al., J Mol Biol, 2009, 10.1016/j.jmb.2009.07.065), XVM2 with 0.6% agar (Wengelnik et al., J Bacteriol, 1996, 10.1128/jb.178.4.1061-1069.1996), and M9 with 0.6% agar. The green arrows point to examples of the outer membrane tubes that can be seen in the images.

**Fig. S2.**
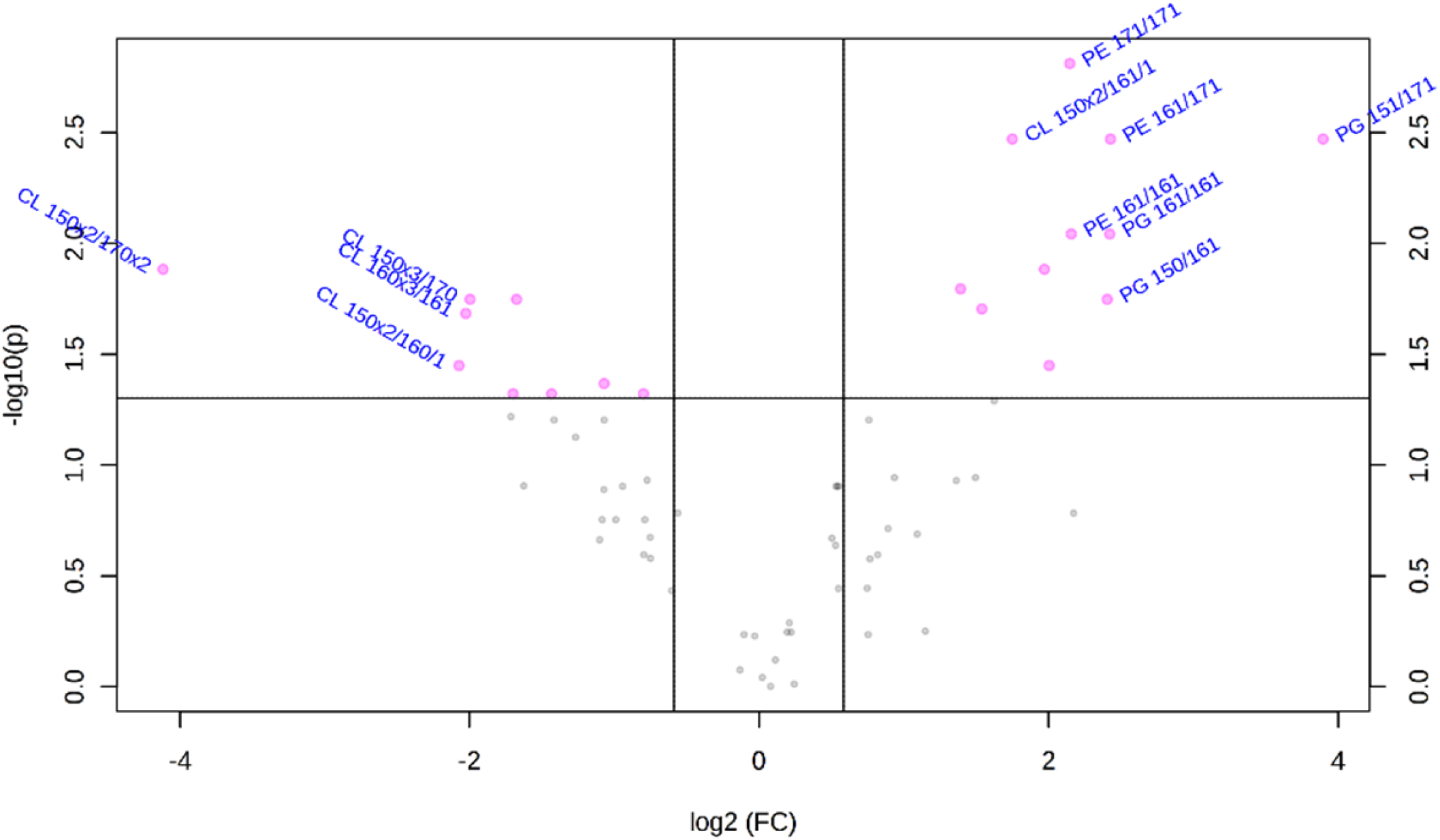
Volcano plot analysis of the lipidomic data. The 20 most altered lipids between the OMV and whole cell samples are identified in the plot as the ones presenting fold change values above 1.5 and p<0.05. Statistical significance was evaluated by FDR-adjusted t-test.

**Fig. S3.**
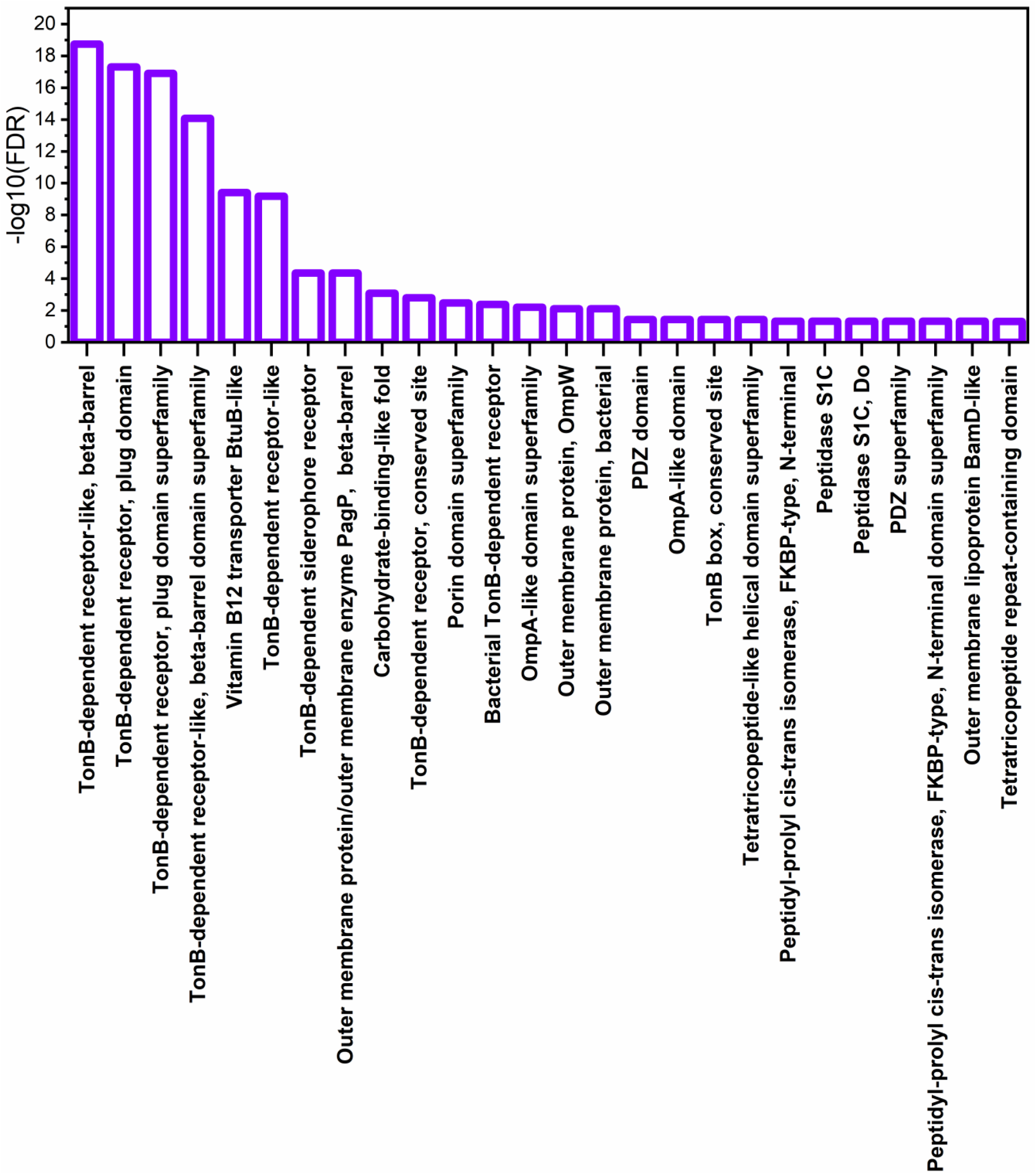
Most significantly enriched InterPro domains found in the purified OMVs compared to the *X. citri* pv. *citri* 306 genome. The lowest false discovery rates (FDR), thus the highest -log10(FDR) values, were observed for domains related to TonB-dependent receptors.

## References

Aebi, C., Stone, B., Beucher, M., Cope, L.D., Maciver, I., Thomas, S.E. et al.. (1996) Expression of the CopB outer membrane protein by Moraxella catarrhalis is regulated by iron and affects iron acquisition from transferrin and lactoferrin. Infect Immun 64: 2024–2030.

Alvarez-Martinez, C.E., Sgro, G.G., Araujo, G.G., Paiva, M.R.N., Matsuyama, B.Y., Guzzo, C.R. et al.. (2021) Secrete or perish: The role of secretion systems in Xanthomonas biology. Computational and Structural Biotechnology Journal 19: 279–302.

Aparna, G., Chatterjee, A., Sonti, R.V., and Sankaranarayanan, R. (2009) A Cell Wall– Degrading Esterase of Xanthomonas oryzae Requires a Unique Substrate Recognition Module for Pathogenesis on Rice. The Plant Cell 21: 1860–1873.

Armenteros, J.J.A., Tsirigos, K.D., Sonderby, C.K., Petersen, T.N., Winther, O., Brunak, S. et al.. (2019) SignalP 5.0 improves signal peptide predictions using deep neural networks. Nature Biotechnology 37: 420-+.

Assis, R.D.B., Polloni, L.C., Patane, J.S.L., Thakur, S., Felestrino, E.B., Diaz-Caballero, J. et al.. (2017) Identification and analysis of seven effector protein families with different adaptive and evolutionary histories in plant-associated members of the Xanthomonadaceae. Scientific Reports 7.

Aznar, A., and Dellagi, A. (2015) New insights into the role of siderophores as triggers of plant immunity: what can we learn from animals? Journal of Experimental Botany 66: 3001–3010.

Bahar, O., Mordukhovich, G., Luu, D.D., Schwessinger, B., Daudi, A., Jehle, A.K. et al.. (2016) Bacterial Outer Membrane Vesicles Induce Plant Immune Responses. Molecular Plant-Microbe Interactions 29: 374–384.

Beltran-Heredia, E., Tsai, F.C., Salinas-Almaguer, S., Cao, F.J., Bassereau, P., and Monroy, F. (2019) Membrane curvature induces cardiolipin sorting. Communications Biology 2.

Biller, S.J., Schubotz, F., Roggensack, S.E., Thompson, A.W., Summons, R.E., and Chisholm, S.W. (2014) Bacterial vesicles in marine ecosystems. Science 343: 183–186.

Blanvillain, S., Meyer, D., Boulanger, A., Lautier, M., Guynet, C., Denance, N. et al.. (2007) Plant Carbohydrate Scavenging through TonB-Dependent Receptors: A Feature Shared by Phytopathogenic and Aquatic Bacteria. Plos One 2.

Bligh, E.G., and Dyer, W.J. (1959) A Rapid Method of Total Lipid Extraction and Purification. Canadian Journal of Biochemistry and Physiology 37: 911–917.

Büttner, D., and Bonas, U. (2010) Regulation and secretion of Xanthomonas virulence factors. Fems Microbiology Reviews 34: 107–133.

Cao, P., and Wall, D. (2019) Direct visualization of a molecular handshake that governs kin recognition and tissue formation in myxobacteria. Nature Communications 10: 3073.

Chaves-Filho, A.B., Pinto, I.F.D., Dantas, L.S., Xavier, A.M., Inague, A., Faria, R.L. et al.. (2019) Alterations in lipid metabolism of spinal cord linked to amyotrophic lateral sclerosis. Scientific Reports 9: 11642.

Coughlin, R.T., Tonsager, S., and Mcgroarty, E.J. (1983) Quantitation of Metal-Cations Bound to Membranes and Extracted Lipopolysaccharide of Escherichia-Coli. Biochemistry 22: 2002–2007.

Dejean, G., Blanvillain-Baufume, S., Boulanger, A., Darrasse, A., de Bernonville, T.D., Girard, A.L. et al.. (2013) The xylan utilization system of the plant pathogen Xanthomonas campestris pv campestris controls epiphytic life and reveals common features with oligotrophic bacteria and animal gut symbionts. New Phytologist 198: 899–915.

Dey, A., and Wall, D. (2014) A Genetic Screen in Myxococcus xanthus Identifies Mutants That Uncouple Outer Membrane Exchange from a Downstream Cellular Response. Journal of Bacteriology 196: 4324–4332.

Dow, J.M., Clarke, B.R., Milligan, D.E., Tang, J.L., and Daniels, M.J. (1990) Extracellular Proteases from Xanthomonas campestris pv. Campestris, the Black Rot Pathogen. Applied and Environmental Microbiology 56: 2994–2998.

Elhenawy, W., Debelyy, M.O., and Feldman, M.F. (2014) Preferential Packing of Acidic Glycosidases and Proteases into Bacteroides Outer Membrane Vesicles. Mbio 5.

Evans, A.G.L., Davey, H.M., Cookson, A., Currinn, H., Cooke-Fox, G., Stanczyk, P.J., and Whitworth, D.E. (2012) Predatory activity of Myxococcus xanthus outer-membrane vesicles and properties of their hydrolase cargo. Microbiology 158: 2742–2752.

Feitosa-Junior, O.R., Stefanello, E., Zaini, P.A., Nascimento, R., Pierry, P.M., Dandekar, A.M. et al.. (2019) Proteomic and Metabolomic Analyses of Xylella fastidiosa OMV-Enriched Fractions Reveal Association with Virulence Factors and Signaling Molecules of the DSF Family. Phytopathology 109: 1344–1353.

Fett, W.F., Gerard, H.C., Moreau, R.A., Osman, S.F., and Jones, L.E. (1992) Screening of Nonfilamentous Bacteria for Production of Cutin-Degrading Enzymes. Applied and Environmental Microbiology 58: 2123–2130.

Figaj, D., Ambroziak, P., Przepiora, T., and Skorko-Glonek, J. (2019) The Role of Proteases in the Virulence of Plant Pathogenic Bacteria. International Journal of Molecular Sciences 20.

Fischer, T., Schorb, M., Reintjes, G., Kolovou, A., Santarella-Mellwig, R., Markert, S. et al.. (2019) Biopearling of Interconnected Outer Membrane Vesicle Chains by a Marine Flavobacterium. Applied and Environmental Microbiology 85: e00829–00819.

Franceschini, A., Szklarczyk, D., Frankild, S., Kuhn, M., Simonovic, M., Roth, A. et al.. (2013) STRING v9.1: protein-protein interaction networks, with increased coverage and integration. Nucleic Acids Research 41: D808–D815.

Guerrero-Mandujano, A., Hernandez-Cortez, C., Ibarra, J.A., and Castro-Escarpulli, G. (2017) The outer membrane vesicles: Secretion system type zero. Traffic 18: 425–432.

Hampton, C.M., Guerrero-Ferreira, R.C., Storms, R.E., Taylor, J.V., Yi, H., Gulig, P.A., and Wright, E.R. (2017) The Opportunistic Pathogen Vibrio vulnificus Produces Outer Membrane Vesicles in a Spatially Distinct Manner Related to Capsular Polysaccharide. Frontiers in Microbiology 8.

Hase, C.C., and Finkelstein, R.A. (1993) Bacterial Extracellular Zinc-Containing Metalloproteases. Microbiological Reviews 57: 823–837.

Hassani, M.A., Duran, P., and Hacquard, S. (2018) Microbial interactions within the plant holobiont. Microbiome 6.

Hellman, J., Loiselle, P.M., Zanzot, E.M., Allaire, J.E., Tehan, M.M., Boyle, L.A. et al.. (2000) Release of gram-negative outer-membrane proteins into human serum and septic rat blood and their interactions with immunoglobulin in antiserum to Escherichia coli J5. J Infect Dis 181: 1034–1043.

Hickey, C.A., Kuhn, K.A., Donermeyer, D.L., Porter, N.T., Jin, C., Cameron, E.A. et al.. (2015) Colitogenic Bacteroides thetaiotaomicron Antigens Access Host Immune Cells in a Sulfatase-Dependent Manner via Outer Membrane Vesicles. Cell Host Microbe 17: 672–680.

Hou, S.G., Jamieson, P., and He, P. (2018) The cloak, dagger, and shield: proteases in plant-pathogen interactions. Biochemical Journal 475: 2491–2509.

Katsir, L., and Bahar, O. (2017) Bacterial outer membrane vesicles at the plant-pathogen interface. Plos Pathogens 13.

Kim, Y.R., Kim, B.U., Kim, S.Y., Kim, C.M., Na, H.S., Koh, J.T. et al.. (2010) Outer membrane vesicles of Vibrio vulnificus deliver cytolysin-hemolysin Vvha into epithelial cells to induce cytotoxicity. Biochemical and Biophysical Research Communications 399: 607–612.

Krewulak, K.D., and Vogel, H.J. (2011) TonB or not TonB: is that the question? Biochemistry and Cell Biology 89: 87–97.

Lappann, M., Otto, A., Becher, D., and Vogel, U. (2013) Comparative proteome analysis of spontaneous outer membrane vesicles and purified outer membranes of Neisseria meningitidis. J Bacteriol 195: 4425–4435.

Lin, T.Y., and Weibel, D.B. (2016) Organization and function of anionic phospholipids in bacteria. Applied Microbiology and Biotechnology 100: 4255–4267.

Louden, B.C., Haarmann, D., and Lynne, A.M. (2011) Use of Blue Agar CAS Assay for Siderophore Detection. Journal of Microbiology &amp; Biology Education 12: 51–53.

Macdonald, I.A., and Kuehn, M.J. (2013) Stress-induced outer membrane vesicle production by Pseudomonas aeruginosa. J Bacteriol 195: 2971–2981.

Maredia, R., Devineni, N., Lentz, P., Dallo, S.F., Yu, J., Guentzel, N. et al.. (2012) Vesiculation from Pseudomonas aeruginosa under SOS. ScientificWorldJournal 2012: 402919.

McBroom, A.J., and Kuehn, M.J. (2007) Release of outer membrane vesicles by Gram-negative bacteria is a novel envelope stress response. Mol Microbiol 63: 545–558.

McCaig, W.D., Koller, A., and Thanassi, D.G. (2013) Production of Outer Membrane Vesicles and Outer Membrane Tubes by Francisella novicida. Journal of Bacteriology 195: 1120–1132.

Nascimento, R., Gouran, H., Chakraborty, S., Gillespie, H.W., Almeida-Souza, H.O., Tu, A. et al.. (2016) The Type II Secreted Lipase/Esterase LesA is a Key Virulence Factor Required for Xylella fastidiosa Pathogenesis in Grapevines. Scientific Reports 6.

Ou, S.H. (1985) Bacterial leaf blight. In Rice Diseases: Commonwealth Mycological Institute, pp. 61–96.

Pirbadian, S., Barchinger, S.E., Leung, K.M., Byun, H.S., Jangir, Y., Bouhenni, R.A. et al.. (2014) Shewanella oneidensis MR-1 nanowires are outer membrane and periplasmic extensions of the extracellular electron transport components. Proceedings of the National Academy of Sciences of the United States of America 111: 12883–12888.

Pirbadian, S., Barchinger, S.E., Leung, K.M., Byun, H.S., Jangir, Y., Bouhenni, R.A. et al.. (2015) Bacterial Nanowires of Shewanella Oneidensis MR-1 are Outer Membrane and Periplasmic Extensions of the Extracellular Electron Transport Components. Biophysical Journal 108: 368a–368a.

Planas-Iglesias, J., Dwarakanath, H., Mohammadyani, D., Yanamala, N., Kagan, V.E., and Klein-Seetharaman, J. (2015) Cardiolipin Interactions with Proteins. Biophysical Journal 109: 1282–1294.

Rakoff-Nahoum, S., Coyne, M.J., and Comstock, L.E. (2014) An ecological network of polysaccharide utilization among human intestinal symbionts. Curr Biol 24: 40–49.

Ramnath, L., Sithole, B., and Govinden, R. (2017) Identification of lipolytic enzymes isolated from bacteria indigenous to Eucalyptus wood species for application in the pulping industry. Biotechnology Reports 15: 114–124.

Remis, J.P., Wei, D.G., Gorur, A., Zemla, M., Haraga, J., Allen, S. et al.. (2014) Bacterial social networks: structure and composition of Myxococcus xanthus outer membrane vesicle chains. Environmental Microbiology 16: 598–610.

Renner, L.D., and Weibel, D.B. (2011) Cardiolipin microdomains localize to negatively curved regions of Escherichia coli membranes. Proceedings of the National Academy of Sciences of the United States of America 108: 6264–6269.

Sampath, V., McCaig, W.D., and Thanassi, D.G. (2018) Amino acid deprivation and central carbon metabolism regulate the production of outer membrane vesicles and tubes by Francisella. Molecular Microbiology 107: 523–541.

Schlechter, R.O., Miebach, M., and Remus-Emsermann, M.N.P. (2019) Driving factors of epiphytic bacterial communities: A review. Journal of Advanced Research 19: 57–65.

Schwechheimer, C., and Kuehn, M.J. (2013) Synthetic effect between envelope stress and lack of outer membrane vesicle production in Escherichia coli. J Bacteriol 195: 4161–4173.

Schwechheimer, C., and Kuehn, M.J. (2015) Outer-membrane vesicles from Gram-negative bacteria: biogenesis and functions. Nature Reviews Microbiology 13: 605–619.

Schwyn, B., and Neilands, J.B. (1987) Universal Chemical-Assay for the Detection and Determination of Siderophores. Analytical Biochemistry 160: 47–56.

Sgro, G.G., Oka, G.U., Souza, D.P., Cenens, W., Bayer-Santos, E., Matsuyama, B.Y. et al.. (2019) Bacteria-Killing Type IV Secretion Systems. Frontiers in Microbiology 10.

Shetty, A., Chen, S.C., Tocheva, E.I., Jensen, G.J., and Hickey, W.J. (2011) Nanopods: A New Bacterial Structure and Mechanism for Deployment of Outer Membrane Vesicles. Plos One 6.

Sidhu, V.K., Vorholter, F.J., Niehaus, K., and Watt, S.A. (2008) Analysis of outer membrane vesicle associated proteins isolated from the plant pathogenic bacterium Xanthomonas campestris pv. campestris. Bmc Microbiology 8.

Sjöström, A.E., Sandblad, L., Uhlin, B.E., and Wai, S.N. (2015) Membrane vesicle-mediated release of bacterial RNA. Scientific Reports 5.

Solé, M., Scheibner, F., Hoffmeister, A.K., Hartmann, N., Hause, G., Rother, A. et al.. (2015) Xanthomonas campestris pv. vesicatoria Secretes Proteases and Xylanases via the Xps Type II Secretion System and Outer Membrane Vesicles. Journal of Bacteriology 197: 2879–2893.

Sorice, M., Manganelli, V., Matarrese, P., Tinari, A., Misasi, R., Malorni, W., and Garofalo, T. (2009) Cardiolipin-enriched raft-like microdomains are essential activating platforms for apoptotic signals on mitochondria. Febs Letters 583: 2447–2450.

Tamir-Ariel, D., Rosenberg, T., Navon, N., and Burdman, S. (2012) A secreted lipolytic enzyme from Xanthomonas campestris pv. vesicatoria is expressed in planta and contributes to its virulence. Molecular Plant Pathology 13: 556–567.

Tashiro, Y., Inagaki, A., Shimizu, M., Ichikawa, S., Takaya, N., Nakajima-Kambe, T. et al.. (2011) Characterization of Phospholipids in Membrane Vesicles Derived from Pseudomonas aeruginosa. Bioscience Biotechnology and Biochemistry 75: 605–607.

Tayi, L., Maku, R.V., Patel, H.K., and Sonti, R.V. (2016) Identification of Pectin Degrading Enzymes Secreted by Xanthomonas oryzae pv. oryzae and Determination of Their Role in Virulence on Rice. Plos One 11.

Toledo, A., Coleman, J.L., Kuhlow, C.J., Crowley, J.T., and Benach, J.L. (2012) The enolase of Borrelia burgdorferi is a plasminogen receptor released in outer membrane vesicles. Infect Immun 80: 359–368.

Toyofuku, M., Nomura, N., and Eberl, L. (2019) Types and origins of bacterial membrane vesicles. Nature Reviews Microbiology 17: 13–24.

Turner, P., Barber, C., and Daniels, M. (1984) Behavior of the Transposons Tn5 and Tn7 in Xanthomonas-Campestris Pv Campestris. Molecular & General Genetics 195: 101–107.

Tyanova, S., Temu, T., and Cox, J. (2016) The MaxQuant computational platform for mass spectrometry-based shotgun proteomics. Nature Protocols 11: 2301–2319.

Ueda, H., Kurose, D., Kugimiya, S., Mitsuhara, I., Yoshida, S., Tabata, J. et al.. (2018) Disease severity enhancement by an esterase from non-phytopathogenic yeast Pseudozyma antarctica and its potential as adjuvant for biocontrol agents. Scientific Reports 8.

Yu, N.Y., Wagner, J.R., Laird, M.R., Melli, G., Rey, S., Lo, R. et al.. (2010) PSORTb 3.0: improved protein subcellular localization prediction with refined localization subcategories and predictive capabilities for all prokaryotes. Bioinformatics 26: 1608–1615.

Zwarycz, A.S., Livingstone, P.G., and Whitworth, D.E. (2020) Within-species variation in OMV cargo proteins: the Myxococcus xanthus OMV pan-proteome. Molecular Omics 16: 387–397.

